# Magnetic field-induced ER stress reprograms the tumor microenvironment to improve triple-negative breast cancer survival

**DOI:** 10.64898/2026.03.22.713285

**Authors:** Vivek Sharma, Chandra Khantwal, Kishori Konwar

**Affiliations:** Asha Medical; Murigenics; ScellAI, LLC

## Abstract

**Background:** Non-invasive electromagnetic field (EMF)-based therapies offer a potential route to modulate local tumor–immune interactions but their mechanistic basis remains poorly defined.

**Methods:** We evaluated Asha therapy, a proprietary low-intensity (50khz, 2 mT, 25% duty cycle) alternating magnetic-field treatment in preclinical breast cancer models. Cellular responses in human triple negative breast cancer cell lines (MDA-MB-231 and MDA-MB-468) were evaluated using bulk RNA sequencing, quantitative proteomics, flow cytometry, and cytokine analysis and proteomics analysis. Tumor microenvironment responses in mouse 4T1 breast cancer model was characterized using single-cell CITE-seq analysis. Functional efficacy was assessed in vivo using the murine 4T1 triple-negative breast cancer model, both as monotherapy and in combination with anti-PD1 checkpoint blockade. Clinical relevance was assessed by deriving a 19-gene neutrophil activation signature from Asha-induced transcriptional changes and projecting it onto two independent TNBC patient cohorts (METABRIC n=338, SCAN-B n=874) for survival analysis.

**Results:** Asha therapy induced endoplasmic reticulum (ER) stress and activated an adaptive unfolded-protein response in tumor cells, triggering robust NF-κB and interferon signaling and time-dependent secretion of inflammatory cytokines. In vivo, these tumor-intrinsic changes propagated to the tumor microenvironment (TME), reprogramming fibroblasts from contractile states to immune-recruiting, interferon-responsive phenotypes and enriching for interferon-stimulated, metabolically active neutrophils and macrophages. These coordinated innate immune changes occurred without overt cytotoxicity and were associated with significant reductions in metastasis and improved survival. Combination with anti-PD1 therapy markedly enhanced efficacy, reducing lung metastasis and mortality by 88% compared with control. The neutrophil activation signature derived from Asha-treated tumors was associated with improved overall survival in both METABRIC (log-rank p=0.036) and SCAN-B (p=0.048) TNBC cohorts by Kaplan-Meier analysis, with pooled multivariable Cox regression confirming significant survival benefit (HR=0.75, 95% CI 0.59-0.94, p=0.01).

**Conclusions:** Asha therapy triggers a controlled ER stress response in tumor cells that drives interferon-mediated cytokine release and immune reprogramming of the TME, resulting in anti-metastatic and survival benefits. These findings identify electromagnetic-field exposure as a potential non-pharmacologic strategy to activate innate immunity and sensitize tumors to checkpoint blockade, supporting further clinical development of EMF-based immunotherapy.

## Introduction

Cancer immunotherapy has transformed oncology by enabling durable tumor control through immune modulation^1^. Immune checkpoint inhibitors targeting PD-1, PD-L1, and CTLA-4 have produced lasting responses in several malignancies, yet only a subset of patients benefit^2–9^. Across most solid tumors, objective response rates remain 10–40%, and treatment is often limited by immune-related toxicities, with grade ≥3 events occurring in up to half of patients^10–13^. Overcoming primary or acquired resistance while minimizing systemic toxicity remains a central challenge.

Triple-negative breast cancer (TNBC) exemplifies these challenges. Although pembrolizumab combined with chemotherapy improves outcomes in PD-L1+ TNBC, the majority of patients show minimal benefit with only 2-3 month progression free survival (PFS) gains, and treatment-related grade ≥3 toxicity occurs in nearly 30% of patients^9,14^. Furthermore, patients with visceral and central nervous system (CNS) metastases that are common in advanced TNBC were underrepresented or excluded from pivotal clinical trials, and retrospective data suggest poor immunotherapy responses in these sites^15,16^.

A key determinant of immunotherapy success is the modulation of the tumor microenvironment (TME)^17^. In TNBC, checkpoint resistance is multifactorial, involving not only limited T cell infiltration but also accumulation of immunosuppressive myeloid populations and regulatory T cells, as well as the formation of stromal barriers that exclude or functionally impair effector lymphocytes^18,19^ Numerous strategies have sought to reprogram the immunosuppressive TME, including radiation, oncolytic viruses, and targeted cytokine delivery, but these interventions often introduce additional toxicity or logistical complexity^20–22^. Recent evidence indicates that myeloid cell reprogramming, particularly conversion of neutrophils from immunosuppressive to anti-tumor phenotypes, may be critical for checkpoint blockade efficacy^23,24^.

Non-invasive physical modalities such as electromagnetic fields (EMF) represent a promising but underexplored means of immune activation. For example, Novocure’s Tumor Treating Fields (TTFields) platform delivers low-intensity, intermediate-frequency electric fields and is clinically approved for glioblastoma, malignant pleural mesothelioma, and metastatic non-small cell lung cancer^25–27^. Recent preclinical studies indicate that TTFields can induce immunogenic cell death and enhance antigen presentation, and TTFields is being evaluated in combination with checkpoint blockade therapy across several tumor types^28,29^. Similarly, the Houston Methodist team’s Oncomagnetic device applies oscillating EMF to disrupt mitochondrial metabolism and elevate reactive oxygen species selectively in cancer cells, with early evidence of downstream immune effects in gliomas^30,31^. Most recently, it was demonstrated that low-intensity induced EMF reduce TNBC tumor growth and metastasis through immune modulation^32^. Together, these studies establish that EMF exposure can engage tumor-immune interactions, yet the molecular mechanisms initiating TME remodeling remain undefined, limiting rational therapeutic development.

The present study was motivated by the disocvery that cancer cells exposed to Asha therapy, a low-intensity (2 mT) 50 kHz alternating magnetic-field treatment, secreted inflammatory cytokines despite minimal direct cytotoxicity. This disconnect between robust molecular reprogramming and preserved cell viability suggests an adaptive stress response rather than damage-associated molecular patterns. Thus, we hypothesized that ER stress acts as the main molecular bridge between EMF exposure and the immune response. This idea is based on three key observations: EMF can trigger stress pathways in mammalian cells,^33–35^ protein-folding stress is known to drive inflammatory signaling,^36^ and these resulting ‘danger signals’ have the potential to flip a suppressive TME into one that actively fights the cancer.^37^

Using integrated transcriptomic, proteomic, and single-cell analyses, we found that Asha therapy triggers a controlled ER stress in cancer cells. This initiates a pro-inflammatory immune activation characterized by the enrichment of interferon-stimulated neutrophils, macrophage repolarization, and enhanced T cell priming. Critically, this reprogramming sensitizes tumors to PD-1 blockade, reducing metastatic burden by over 90% and mortality by 88% in combination therapy. These findings position ER stress as the mechanistic foundation for EMF-induced immunomodulation, provide a framework for optimizing EMF parameters, and inspire potential therapeutic combinations with diverse immunotherapies.

## Methods

### Cell culture

MDA-MB-231 cells were cultured in RPMI-1640 medium (ThermoFisher #11835030) supplemented with 10% fetal bovine serum (FBS, ThermoFisher #A5209501) and 1% antibiotic-antimycotic solution (ThermoFisher #15240062). MDA-MB-468 cells were cultured in Dulbecco’s Modified Eagle Medium (DMEM, ThermoFisher # 11960044) supplemented with 10% FBS and 1% Antibiotic-Antimycotic solution. MCF-10A cells were cultured in specialized media (Elab Biosciences # CM-0525) supplemented with 10% FBS and 1% Antibiotic-Antimycotic solution. All cells were maintained at 37°C in a humidified atmosphere with 5% CO₂. Cell media was replaced every 2-3 days, and cells were typically passaged once they reached 50-80% confluence, and used within 20 passages

### Asha therapy exposure

For in vitro experiments, Asha therapy (50 kHz, 2 mT, sinusoidal waveform, 25% duty cycle) was delivered using a 75-mm radius Helmholtz coil placed inside a dedicated cell-culture incubator. The coil was designed to accommodate standard culture plates. Finite-element electromagnetic modeling (COMSOL Multiphysics) confirmed uniform EMF variation across the plate area. Temperature was monitored using a fiber-optic thermocouple (Osensa PRB-100-1M-STM-MRI) placed in direct contact with the plate, and the incubator set point was adjusted to maintain 37.0 ± 0.5 °C throughout the experiment. A calibrated search coil was used to monitor field amplitude at the plate location. Therapy was delivered continuously for the full duration of each experiment. As the EMF would contaminate nearby plates, control samples were maintained in a separate but otherwise identical incubator. Environmental conditions (temperature, humidity, CO₂, and plate format) were matched across both incubators, and control plates were handled in parallel with therapy plates to ensure identical timing of media changes, passaging, and sampling.

For in vivo experiments, Asha therapy was delivered using a custom 185-mm radius Helmholtz coil designed to house a standard mouse cage containing 5–6 mice. The coil generated a uniform electromagnetic field that provided full-body exposure to all animals in the cage. A calibrated search coil was used to measure the magnetic field amplitude at the cage center to verify correct field delivery (±10% of the target field). Therapy was applied continuously for the duration of the experiment unless otherwise specified.

### Thapsigargin

Thapsigargin (MedChemExpress #HY-13433) stock solution (500 μM) was prepared in DMSO (ThermoFisher # D12345) and stored at −80°C. Working concentrations were prepared weekly by diluting the stock solution in culture medium to achieve the desired final concentration (IC_50_ for 72-hour exposure on MDA-MB-468 cells was previously determined to be 5nM).

### In Vitro Asha Therapy Treatment

For Bulk RNA analysis, flow cytometery and cytokine measurements, 50,000 cells per well were seeded in 0.5 mL of culture medium in 24-well plates and incubated overnight at 37°C. The following morning, thapsigargin (5 nM) was added to designated treatment groups as a positive control for ER stress induction. Plates were then placed either in a standard incubator (control and thapsigargin groups) or in the incubator equipped with the Asha therapy setup for the designated therapy duration.

### Bulk RNA sequencing

Following experimental treatments, culture supernatants were removed and cells were lysed using the Zymo Research RNA extraction reagent according to the manufacturer’s protocol. Total RNA was extracted and shipped to Novogene (Sacramento, CA, USA) for quality control, library preparation, and sequencing. RNA integrity was assessed using an Agilent 2100 Bioanalyzer.

Messenger RNA was purified using poly-T oligo-attached magnetic beads and fragmented prior to library construction. Libraries were prepared using a standard unstranded mRNA-seq workflow with random hexamer priming for first-strand cDNA synthesis, followed by second-strand synthesis (without dUTP marking), end repair, A-tailing, adapter ligation, size selection, and PCR amplification. Sequencing was performed on an Illumina NovaSeq X Plus platform to generate 2×150 bp paired-end reads.

### Bulk RNA sequencing Analysis

Raw paired-end FASTQ files were quality-filtered and adapter-trimmed using fastp (v0.23.2)^38^. Cleaned reads were aligned to the human reference genome GRCh38 using STAR (v2.7.10a) with --quantMode GeneCounts and Ensembl gene annotations (release 113). Strandedness was evaluated from the STAR ReadsPerGene.out.tab files by comparing forward/reverse read ratios (mean = 1.06), confirming an unstranded library type; therefore, read counts were extracted from column 2 (unstranded) of each output file^39^. Differential expression analysis was conducted using PyDESeq2 (version 0.5.2) with default parameters^40,41^. Genes with adjusted p < 0.05 (Benjamini–Hochberg correction), |log₂ fold change| > 0.58, and baseMean > 10 were considered significantly differentially expressed.

For gene set enrichment analysis (GSEA) (GSEApy version 1.0.4), we started from PyDESeq2 outputs and ranked genes by the Wald statistic (t-like score)^42^. Prior to ranking, we applied the following filters/normalization: (i) excluded genes with baseMean < 10; (ii) excluded rows lacking a valid HGNC gene symbol; (iii) collapsed duplicate symbols (multiple Ensembl IDs mapping to the same symbol) by retaining the entry with the largest |Wald statistic|; and (iv) removed rows with missing Wald statistics. Symbols were standardized case-insensitively to ensure uniqueness. GSEA was run against MSigDB Hallmark 2020, KEGG 2021 (Human), Reactome 2022, and GO Biological Process, Cellular Components, and Molecular Function gene sets, using the 15–500 gene set-size filter^43^. Pathways with FDR < 0.05 were considered significant; positive/negative enrichment corresponded to higher/lower expression in Asha vs. control.

### Flow cytometry

Following treatment exposure, cells were detached using trypsin-EDTA (Sigma Aldrich #59428C-500ML) and processed for flow cytometry analysis. Fc receptor blocking was performed using Fc receptor binding inhibitor antibody (ThermoFisher #14-9161-73) according to manufacturer’s instructions. Cell viability was assessed using fixable viability dye (ThermoFisher #65-0866-14). Cells were then fixed and permeabilized using fixation/permeability kit per manufacturer’s protocol (BD Biosciences Cytofix/Cytoperm™ Kit #554715).

Intracellular staining was performed using the following primary antibodies: GRP78-Alexa Fluor 647 (ThermoFisher #CL647-66574), CHOP-PE (Santa Cruz Biotechnology #sc-56107 PE), ATF4 (Santa Cruz Biotechnology #sc-390063 AF594), and xBP1s-Alexa Fluor 488 (ProteinTech #CL488-24868). Flow cytometry acquisition was performed on an Invitrogen Attune NXT flow cytometer, and data were analyzed using De Novo software (FCS Express). Single-stain controls were prepared for each fluorophore and used to calculate compensation matrices in FCS Express. Unstained cells served as negative controls. Flowcytometry gating is detailed in supplementary Figure S1.

### Cell viability assay

MDA-MB-468 cells (5,000 cells in 200 μL media) were plated overnight in black 96-well cell culture plates. The following day, media was replaced with fresh media, and the plates were then placed either in a standard incubator (untreated control group) or in the incubator equipped with the Asha therapy setup for the designated therapy duration. After therapy exposure, cell viability was measured using PrestoBlue™ Cell Viability Reagent (ThermoFisher #A13261) per the manufacturer’s instructions. Briefly, the cell media was removed from each well and replaced with 100 μL of cell media containing 10% PrestoBlue reagent. Two wells on each plate containing media with 10% PrestoBlue reagent but no cells served as blanks for background subtraction. The plates were then placed in a standard incubator for 10 minutes, and the fluorescence was measured on a Thermofisher Varioskan LUX plate reader using an excitation wavelength of 560 nm and an emission wavelength of 590 nm. Fluorescence values were background-subtracted using the blank wells.

### Cytokine analysis

Culture supernatants were collected from MDA-MB-468 cells at 24, 48, 72, and 96 hours following Asha therapy exposure, immediately frozen, and stored at −80 °C until analysis. Samples were shipped on dry ice to Nomic Bio (Montreal, QC, Canada) for quantitative profiling using the nELISA platform, which measured 270 cytokines simultaneously with high sensitivity and multiplex precision^44^. Asha-treated samples were compared with matched control samples at each time point using the non-parametric Mann–Whitney test, and cytokines with a false discovery rate (FDR) < 0.05 were considered significantly altered.

### Proteomic analysis

Following experimental treatments, cells were washed with PBS, pelleted, and shipped on dry ice to PTM Bio (Alameda, CA, USA) for quantitative proteomic analysis. Proteins were extracted in SDS lysis buffer (8 M urea, 5% SDS, protease inhibitors, 50 mM TEAB, pH 8.0), quantified by BCA assay, reduced, alkylated, and digested using the Suspension Trapping (S-Trap) protocol. Peptides were analyzed by LC-MS/MS in Data-Independent Acquisition (DIA) mode on a Thermo Orbitrap Astral mass spectrometer coupled to a Vanquish Neo UHPLC system with a 15 cm × 100 µm C18 column. Samples were separated using a 23-minute gradient (4-99% acetonitrile with 0.1% formic acid) at 0.4-0.9 µL/min. Full MS scans were acquired at 240,000 resolution (m/z 380-980, AGC 500%, 3 ms max IT), with DIA MS/MS using 2 m/z isolation windows (m/z 150-2000, 25% collision energy). Raw data were processed using DIA-NN (v1.9) against the UniProt human reference proteome. Peptide and protein identifications were filtered at 1% false discovery rate (FDR), with label-free quantification based on DIA peak intensities.

Differential protein abundance was analyzed using DESeq2 (via pyDESeq2), which has been validated for mass spectrometry-based proteomics and handles the discrete count-like nature of peptide identification while accounting for technical and biological variability^45^. Analysis was restricted to proteins detected in ≥2 samples per group to ensure reliable quantification. Proteins with padj<0.05 (Benjamini-Hochberg FDR correction) and |log2FC|>0.58 were considered significantly altered. GSEA was performed on reliably quantified proteins (≥2 samples per condition, n=8,534) ranked by sign(log₂FC) × −log₁₀(P-value), using MSigDB Hallmark, Gene Ontology (GO) and KEGG gene sets (FDR<0.05).

### 4T1 lung metastasis study

Female BALB/c mice (5-6 weeks old) were inoculated orthotopically with 500,000 4T1 tumor cells in the mammary fat pad, to model aggressive metastatic disease. Mice were housed in the Murigenics’ (Vallejo, CA) animal facility under standard conditions (12-hour light/dark cycle, 22 ± 2°C) with Laboratory Rodent Diet and municipal tap water provided *ad libitum*. Tumor establishment was confirmed at 7 days post-inoculation by palpable tumor formation (average volume ∼100mm³), and mice were randomized into four groups (n=6 per group) based on tumor size to ensure comparable baseline volumes between groups (Supplementary Figure S2A): isotype control, Asha monotherapy, anti-PD-1 monotherapy, and combination therapy. Mice were weighed twice weekly, and tumor measurements were performed by a trained technician using a digital caliper. Tumor volume was calculated using the formula (length * width * height)/2.

Asha therapy was delivered continuously for the study duration. Isotype control antibody (250 μg, InVivoMAb rat IgG2a isotype control) or anti-PD-1 antibody (250 μg, BioXcell RMP1-14) was administered intraperitoneally twice weekly for a total of 8 doses. At day 28 post-randomization, mice were euthanized via CO₂ inhalation followed by cervical dislocation. Lungs were harvested, and metastatic nodules were quantified by a trained technician blinded to the group assignment.

Statistical analysis was conducted using Kruskal-Wallis test followed by Dunn’s multiple comparisons test. Statistical comparisons were limited to hypothesis-relevant contrasts (monotherapy vs control, combination vs control, and combination vs monotherapy) to maintain statistical power.

### 4T1 survival study

Female BALB/c mice (5-6 weeks old) were inoculated orthotopically with 500,000 4T1 tumor cells in the mammary fat pad. Tumor establishment was confirmed at 7 days post-inoculation by palpable tumor formation (average volume ∼100mm³), and mice were randomized into three groups based on tumor size to ensure comparable baseline volumes between groups: control (n=12), anti-PD-1 monotherapy (n=12), and combination of Asha therapy and anti-PD1 (n=18) (Supplementary Figure S2B): Asha monotherapy was not included as prior experiments had demonstrated superior results with combination therapy. Asha therapy was delivered continuously for the study duration, and anti-PD-1 antibody (250 μg, BioXcell RMP1-14) was administered intraperitoneally twice weekly for a total of 8 doses. 8 days after starting therapy, the primary tumor was resected under anesthesia to model post-surgical metastatic disease, and therapies were paused for a day to allow recovery. Post surgery mice well being was monitored, and their weight was measured twice every week. Mice were sacrificed if their weight decreased or lab staff noticed significant health deterioration. Animal survival was monitored for 107 days after surgery, and the remaining mice were sacrificed at the study end.

### Cellular Indexing of Transcriptomes and Epitopes by sequencing (CITE-seq) on tumor tissue

Female BALB/c mice (6-8 weeks old) were inoculated orthotopically with 100,000 4T1 tumor cells in the mammary fat pad. All animal procedures were approved by Murigenics’ Institutional Animal Care and Use Committee and conducted in accordance with NIH guidelines. Mice were housed in the Murigenics’ (Vallejo, CA) animal facility under standard conditions (12-hour light/dark cycle, 22±2°C) with Laboratory Rodent Diet and municipal tap water provided *ad libitum*. Tumor establishment was confirmed at 10 days post-inoculation by palpable tumor formation (average volume ∼100mm³), and mice were randomized into two groups (n=6 per group) based on tumor size to ensure comparable baseline volumes between groups: untreated control and Asha therapy. Following seven days of treatment, tumor volumes were measured using digital calipers, and mice were euthanized. Tumor tissues were immediately excised, submerged in pre-chilled MACS Tissue Storage Solution (Miltenyi Biotec #130-100-008), and stored at 4°C during transport to MedGenome (Foster City, CA, USA). Three representative tumors from each group (n=3 per condition) were processed for CITE-seq analysis.

Single-cell suspensions were prepared from fresh tumor tissues using enzymatic and mechanical dissociation (MACS mice tumor dissociation kit, Miltenyi Biotec, per manufacturer’s instructions). Cell viability and concentration were assessed, and viable single cells were stained with the TotalSeq™-B Mouse Universal Cocktail (BioLegend) targeting 102 cell surface antigens, including principal immune lineage markers and 7 isotype controls. Cells were then processed for library preparation using the 10x Genomics Chromium Single Cell 3’ v3.1 Dual Index Gene Expression kit according to manufacturer’s protocol^46^. Cells were partitioned into Gel Bead-In-Emulsions (GEMs) for barcoding, followed by reverse transcription, cDNA amplification, and library construction. Paired-end sequencing was performed on an Illumina NovaSeq 6000 platform with a target sequencing depth of approximately 50,000 reads per cell for gene expression (GEX) and 10,000 reads per cell for surface protein (ADT) detection.

Raw sequencing data were processed using Cell Ranger Count v7.1.0 (10x Genomics) for alignment to the mouse reference genome (mm10, 2020-A), barcode processing, and gene expression quantification. Quality control metrics confirmed high-quality data with Phred scores >30 across all read lengths. Preprocessing was performed using CellBender to remove ambient RNA contamination, followed by quality control filtering in Scanpy to retain cells with 200-4000 genes, <10% mitochondrial content, and <15% ribosomal content^47,48^. Doublets were removed using Scrublet, and genes expressed in <10 cells were excluded^49^. Count data were normalized (library size = 10,000) and log-transformed. Integration of RNA and antibody-derived tag (ADT) data with batch correction was performed using TotalVI^50^. UMAP dimensionality reduction and Leiden clustering at resolution 1.0 identified distinct cell populations^51,52^. Cell types were annotated based on RNA canonical marker gene expression and differential expression analysis and confirmed with ADT markers.

Compositional changes were quantified using scCODA, which models cell-type proportions in a Bayesian framework and accounts for the compositional nature of single-cell data, with automatic selection of reference cell type^53^. Statistical credibility was assessed using default FDR threshold of 0.05. Differential expression analysis was performed using the following pseudobulking approach. For gene expression, raw UMI counts (for RNA) were summed per gene across all cells of a given cell type within each biological replicate, yielding a pseudobulk count matrix. DESeq2 (via pyDESeq2) was then applied to these pseudobulk profiles. This approach accounts for both within-sample (technical) and between-sample (biological) variability, avoids pseudoreplication, and provides superior control of false positives compared with single-cell-level Wilcoxon or t-tests^54–56^. For ADT protein measurements, differential abundance was assessed using the Wilcoxon rank-sum test with Benjamini-Hochberg correction for multiple testing. GSEA was performed using GSEApy with MSigDB Hallmark 2020, KEGG 2019 Mouse, Reactome 2022, and Gene Ontology databases (FDR q<0.05).

To assess chromosomal instability in tumor epithelial cells, we calculated copy number variation (CNV) scores based on chromosome-level gene expression variance. Gene chromosomal coordinates were obtained from Ensembl BioMart (Mus musculus GRCm38/mm10, release 102). Genes were mapped to chromosomes 1-19, X, Y, and MT, with 20,740 genes (95.3% of detected genes) successfully annotated with genomic coordinates. To assess chromosomal instability in epithelial cells, we calculated a chromosomal expression heterogeneity score as a proxy for aneuploidy. Gene chromosomal coordinates were obtained from Ensembl BioMart (Mus musculus GRCm38/mm10, release 102) via direct API query. For each cell, mean normalized gene expression was calculated per chromosome for autosomes 1-19 and chromosome X. Chromosomal heterogeneity scores were computed as the coefficient of variation (CV) across chromosomes:

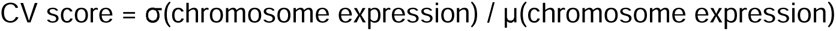

Higher CV scores reflect greater inter-chromosomal expression variance, which correlates with chromosomal copy number alterations in aneuploid cells. While this approach does not resolve specific copy number changes, it provides a continuous metric to distinguish malignant from non-malignant epithelial populations based on chromosomal expression instability patterns^57^.

### Neutrophil activation signature derivation

To construct a human-projectable neutrophil activation signature, differentially expressed genes from Asha-treated tumor-infiltrating neutrophils were ranked by DESeq2 Wald statistic. From the top 100 upregulated transcripts, we excluded ribosomal, mitochondrial, and mouse-specific Gm loci and retained genes mapping to interferon/antiviral pathways, antigen-processing machinery, and treatment-induced myeloid activation. Mouse transcripts without direct human orthologs were replaced with their closest human equivalents (e.g., Gm16337 → IFNL3). This curation yielded a 19-gene “neutrophil activation signature” used for survival analyses. Full details of gene selection, and filtering criteria, are provided in Supplementary S1.

### Human survival analysis (METABRIC cohort)

To test whether our findings translate to human TNBC, we projected our 19-gene neutrophil activation signature onto bulk tumor transcriptomic data from two independent patient cohorts: METABRIC (n=348 patients that has been previously annotated as TNBC^58^, data downloaded from cbioportal.org/study/summary?id=brca_metabric) and SCAN-B (n=874 TNBC patients, data downloaded from data.mendeley.com/datasets/yzxtxn4nmd/4). The neutrophil activation signature was represented by 18 of the 19 candidate genes in both validation cohorts, potentially due to platform specific coverage: *MNDAL* was not availble in the METABRIC microarray data, and *FILIP1L* was not available in the SCAN-B RNA-seq data.

For each patient, we calculated signature scores as the mean normalized expression (z-scores) of the available signature genes. Patients were stratified into high and low signature groups using median split, and overall survival was analyzed using Kaplan-Meier curves and the log-rank test. For each cohort (METABRIC and SCAN-B), we fit Cox proportional hazards models to evaluate the association between high vs low neutrophil activaton signature score and overall survival. Models were adjusted for nodal status (N1+ vs N0), tumor stage (T3+ vs T1–2), and continuous tumor size, with age (>50 vs ≤50 years) included as a stratification factor to account for potential non-proportional hazards.

Meta-analysis across both cohorts was performed using random-effects models to calculate pooled hazard ratios. Hazard ratios (HRs) and 95% confidence intervals (CIs) were reported for the NSG score. All survival analyses were performed using the lifelines Python package (version 0.30.0). This computational approach to project cell-type-specific signatures onto bulk tumor transcriptomes follows established methodologies for myeloid and neutrophil lineage analysis in breast cancer^24,59^.

## Results

### Bulk RNA sequencing reveals therapy induces ER stress and inflammatory response in TNBC cells

To characterize temporal dynamics of transcriptional response, we performed bulk RNA sequencing of MDA-MB-238 cells at 24 and 48 hours post-treatment. Principal component analysis (PCA) revealed minimal transcriptional divergence between Asha-treated and control cells at 24 hours, with clear separation emerging by 48 hours (Figure 1A).

**Figure 1:**
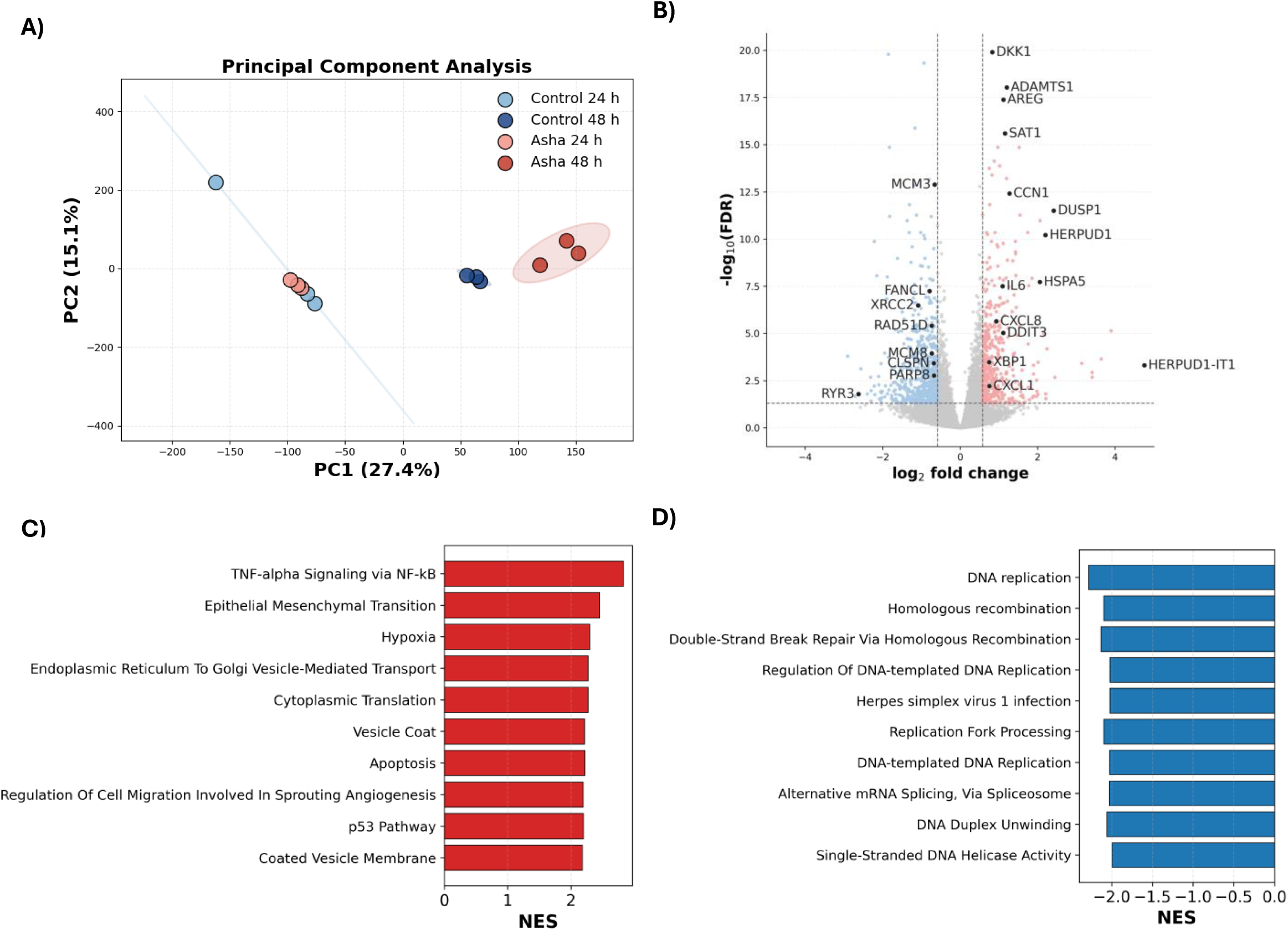
Bulk RNA sequencing reveals progressive transcriptional remodeling and multi-pathway activation following Asha therapy. **A.** Principal Component Analysis (PCA) of normalized gene expression profiles from MDA-MB-231 cells following Asha therapy or control. Each point represents an independent biological replicate (n=3 per condition). Separation along PC1 and PC2 is evident by 48 hours, reflecting robust transcriptional changes at this time point. **B.** Volcano plot displaying the differentially expressed genes (DEGs) in MDA-MB-231 cells following 48hours of Asha therapy compared to control. In this plot, genes were filtered for an adjusted P value {Benjamini–Hochberg corrected) < 0.01, a |log2 fold change| > 0.58. Upregulated genes (red dots) are enriched for stress- and inflammation-associated signaling; downregulated genes (blue dots) are enriched for DNA replication and cell-cycle. **C**. Gene Set Enrichment Analysis (GSEA) of up-regulated pathways in MDA-MB-231 cells at 48 hours, ranked by Normalized Enrichment Score (NES}. GSEA identified 454 significantly up-regulated gene sets (FDR < 0.05), with the most significant enrichment corresponding TNF-alpha signaling via NF-kappaB, indicating activation of inflammatory transcriptional programs. Other major themes include ER-stress/UPR terms, ribosome biogenesis, and mitochondrial metabolism. **D**. GSEA of down-regulated pathways in MDA-MB-231 cells at 48 hours, ranked by NES. GSEA identified 21 significantly down-regulated gene sets (FDR < 0.05). Down-regulated pathways are primarily related to cell-cycle regulation, DNA replication, and DNA repair, consistent with transcriptional repression of replication-associated genes at this time point.

#### Differential gene expression (DGE)

At 48 hours, the significantly (padj<0.05, |log_2_FC|>0.58, baseMean>10; Figure 1B; Supplementary Table S1) upregulated genes included the canonical ER-stress markers, including *HSPA5* (*GRP78*), *DDIT3* (CHOP), and XBP1. Inflammatory mediators including *IL6*, *CXCL8*/*IL8*, and *CXCL1* were also upregulated, as were stress-linked secreted factors including *AREG*, *DKK1*, and *ADAMTS1*. Multiple genes belonging to the DNA replication/repair and cell-cycle control were decreased, including: *XRCC2*, *RAD51D*, and *FANCL*.

### Gene set enrichment analysis

At 48 hours, the most significant (FDR < 0.05) enrichment corresponded to TNF-α signaling via NF-κB (NES = 2.74), indicating activation of inflammatory transcriptional programs. Other up-regulated pathways included multiple ER-stress and unfolded-protein-response terms (Hallmark UPR; GO and KEGG ER-stress modules) mirroring our DGE analysis, as well as ribosome biogenesis and protein synthesis, mitochondrial metabolism, and extracellular-matrix and cytoskeletal remodeling (Figure 1C, Supplementary Table S2). Signficantly down-regulated pathways were primarily related to cell-cycle regulation, DNA replication, and DNA repair, consistent with transcriptional suppression of replication-associated genes at 48 hours (Figure 1D, Supplementary Table S2).

These data demonstrate that Asha therapy induces coordinated ER stress and inflammatory signaling in TNBC cells while suppressing proliferative programs. The concurrent upregulation of ER stress markers and inflammatory cytokines suggests that ER stress triggers danger-associated molecular patterns that could activate immune responses.

### Unbiased global proteomic analysis reveals stress-induced metabolic remodeling in TNBC cells

To complement transcriptomic analysis and capture post-translational regulation, we performed unbiased global proteomic profiling of Asha-treated TNBC cells. Mass spectrometry-based label-free proteomic profiling of Asha-treated versus control MDA-MB-231 cells at 96 hours (selected to capture stable protein-level changes following the 48 hour transcriptional response) identified 8,882 proteins. PCA demonstrated clear separation between control and Asha-treated samples (Fig. 2A), indicating distinct proteomic profiles.

**Figure 2:**
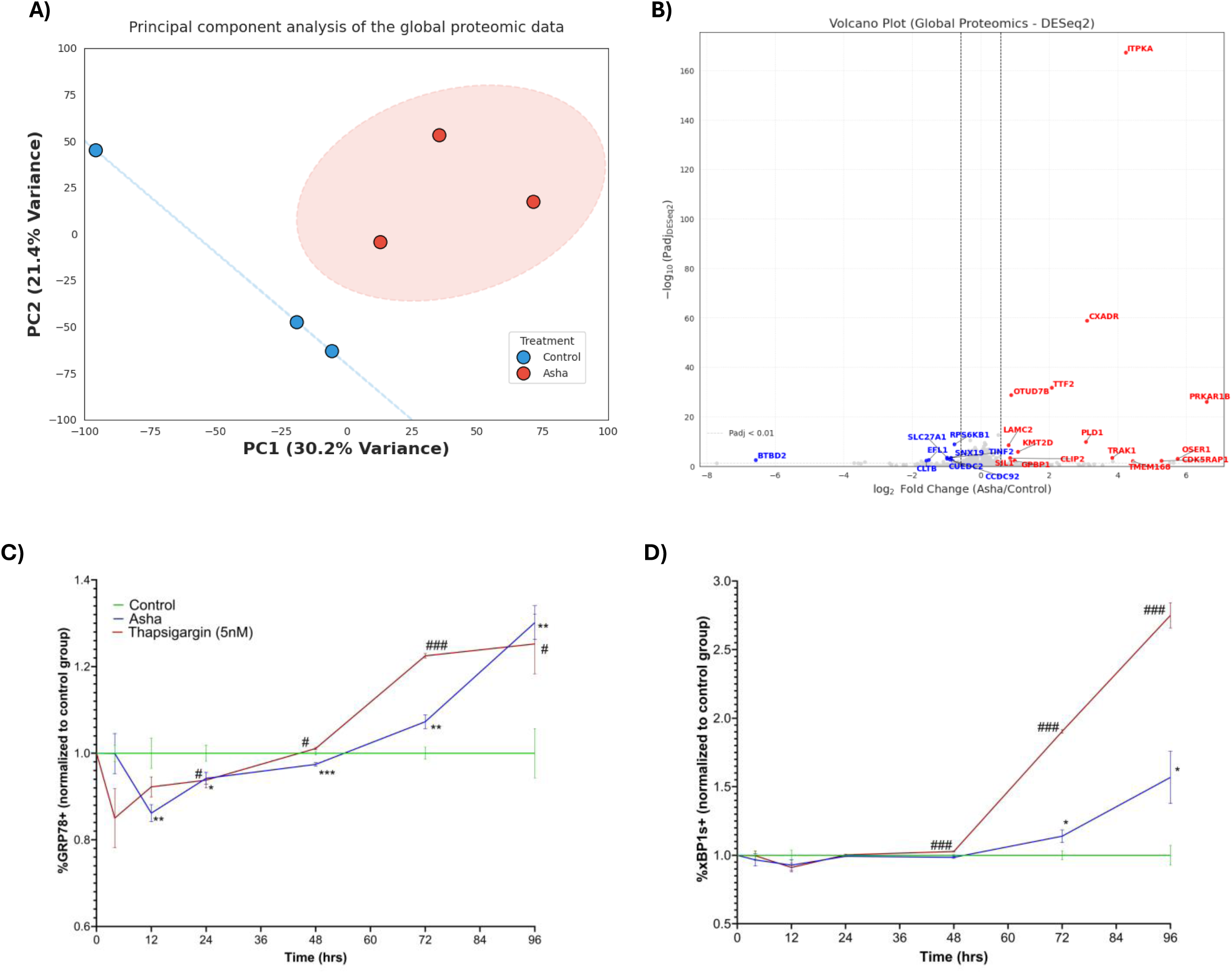
Asha therapy induces ER stress and proteome remodeling in TNBC cells. **(A)** Principal Component Analysis (PCA) of global proteomic profiles from control and Asha-treated MDA-MB-231 cells at 96 hours. Each point represents an independent biological replicate (n=3 per condition). Clear separation indicates distinct proteomic responses to treatment**. (B)** Volcano plot of differentially expressed proteins in MDA-MB-231 cells at 96 hours post-Asha treatment. Analysis performed using label-free quantification and two-sided Student’s t-test. Proteins were filtered for significance based on p < 0.05 and |log2 fold change| > 0.58. A total of 40 proteins were significantly altered (21 upregulated, red dots; 19 downregulated, blue dots). Upregulated proteins are enriched for signaling, proteostasis, and inflammatory components, while downregulated proteins are associated with protein quality control and metabolism. **(C–D)** Time-course analysis of ER stress markers in MDA-MB-468 cells exposed to Asha therapy, thapsigargin (5 nM, positive control), or left untreated (control) for up to 96 hours. Levels of GRP78 (C) and xBP1s (D) were measured by flow cytometry. CHOP and ATF4 showed similar trends with more modest induction (Supplementary Figure S3 A-B). Data are presented as mean ± SEM (n=6 per group per time point), normalized to the control group at each time point. Statistical significance was determined using multiple unpaired t-tests comparing each treatment group to the control group at each time point. Significance markers: *, # (p < 0.05); **, ## (p < 0.01); ***, ### (p < 0.001)

Differential analysis of proteins reliably detected in both conditions (≥2 samples per group) identified 88 significantly altered proteins (padj<0.05, |log2FC|>0.32, Figure 2B, Supplementary Table S3). Key upregulated proteins were primarily associated with ER stress and calcium signaling pathways, including ITPKA (18.9-fold), CXADR (8.6-fold), and PLD1 (8.4-fold). We also observed a massive induction of PRKAR1B (97-fold), a regulatory subunit of protein kinase A (PKA) that mediates stress-adaptive signaling. Conversely, downregulated proteins included the mTOR effector RPS6KB1, the cancer stemness marker ALDH1A3, and DNA maintenance factors such as TINF2 and EFL1. Additionally, 17 proteins were newly detectable following Asha therapy (≥2/3 Asha samples, undetectable in controls), signifying the activation of previously silent transcriptional programs (Supplementary Table S4). These included the master ER-stress regulator NUPR1 and the nutrient-stress sensor ASNSD1.

Together, these findings indicate that Asha therapy inhibits the proliferative and "stem-like" state characteristic of steady-state TNBC tumor cells that is marked by the collapse of mTOR signaling and DNA maintenance, while driving TNBC tumor cells into a stress-adapted secretory profile. By activating master regulators like NUPR1 and the PKA signaling brake (PRKAR1B), Asha-treated tumor cells shift from a mode of rapid growth to one of "intrinsic danger signaling." This proteomic rewiring provides a mechanistic bridge between the initial physical stimulus of the magnetic field and the downstream immune infiltration observed in our in vivo studies, suggesting that the therapy effectively "unmasks" the tumor by triggering a stress program.

### Flow cytometry results indicate Asha therapy induces ER stress and UPR activation in TNBC cancer cells

To characterize the temporal dynamics of ER stress responses, we performed a time-course analysis using the MDA-MB-468 TNBC cell line measuring ER stress marker GRP78, and UPR markers (xBP1s, CHOP and ATF4)^60,61^. We used MDA-MB-468 cells for the time-course experiment because their ER stress markers change over a broader range after stimulation, making it easier to track early and late phases of the UPR. Flow cytometry measurements were obtained 6, 12, 24, 48, 72, and 96 hours after Asha treatment (n=6/group/time point), and thapsigargin (5 nM, 72-hour IC50 value derived experimentally, a SERCA pump inhibitor that depletes ER calcium stores) was used as a positive control for ER stress^62^.

Both Asha and thapsigargin treatment groups exhibited an initial decrease in GRP78, ATF4, xBP1s, and CHOP during the first 12-48 hours, followed by gradual recovery and elevation between 72-96 hours (Figure 2C-2D, Supplementary Figure S3A-3B). Notably, thapsigargin induced robust increases in all UPR markers by 96 hours, while Asha therapy produced more modest elevations (significant in GRP78, xBP1s and CHOP), without reducing cell viability (Supplementary Figure S3C). These results demonstrate adaptive ER stress, a controlled stress response that protects cells rather than killing them, in contrast to the toxic stress induced by high-dose thapsigargin. This pattern matches our transcriptomic and proteomic data showing stress-adaptive signaling without activation of cell death pathways.

### Cytokine analysis of the cell supernatant indicate Asha therapy induces release of pro-inflammatory cytokines by TNBC cancer cells

ER stress is known to trigger secretion of pro-inflammatory cytokines and damage-associated molecular patterns that can activate immune responses^36^. To characterize how Asha-induced ER stress translates into altered tumor secretory programs, we profiled 270 cytokines in conditioned media from MDA-MB-468 cells at 24, 48, 72, and 96 hours using the nELISA platform. Conditioned media showed temporal shifts in secreted factors (Fig. 3A-D). Early responses (24h) included matrix remodeling enzymes (uPA, ADAMTS1), followed by transient inflammatory signals at 48h (IL-6, Tissue Factor). By 72-96h, a sustained immune-recruiting program emerged, with increased secretion of chemokines with known roles in myeloid and lymphoid cell recruiment such as CCL2, CXCL10, CCL20, and CXCL1^63–65^.

**Figure 3.**
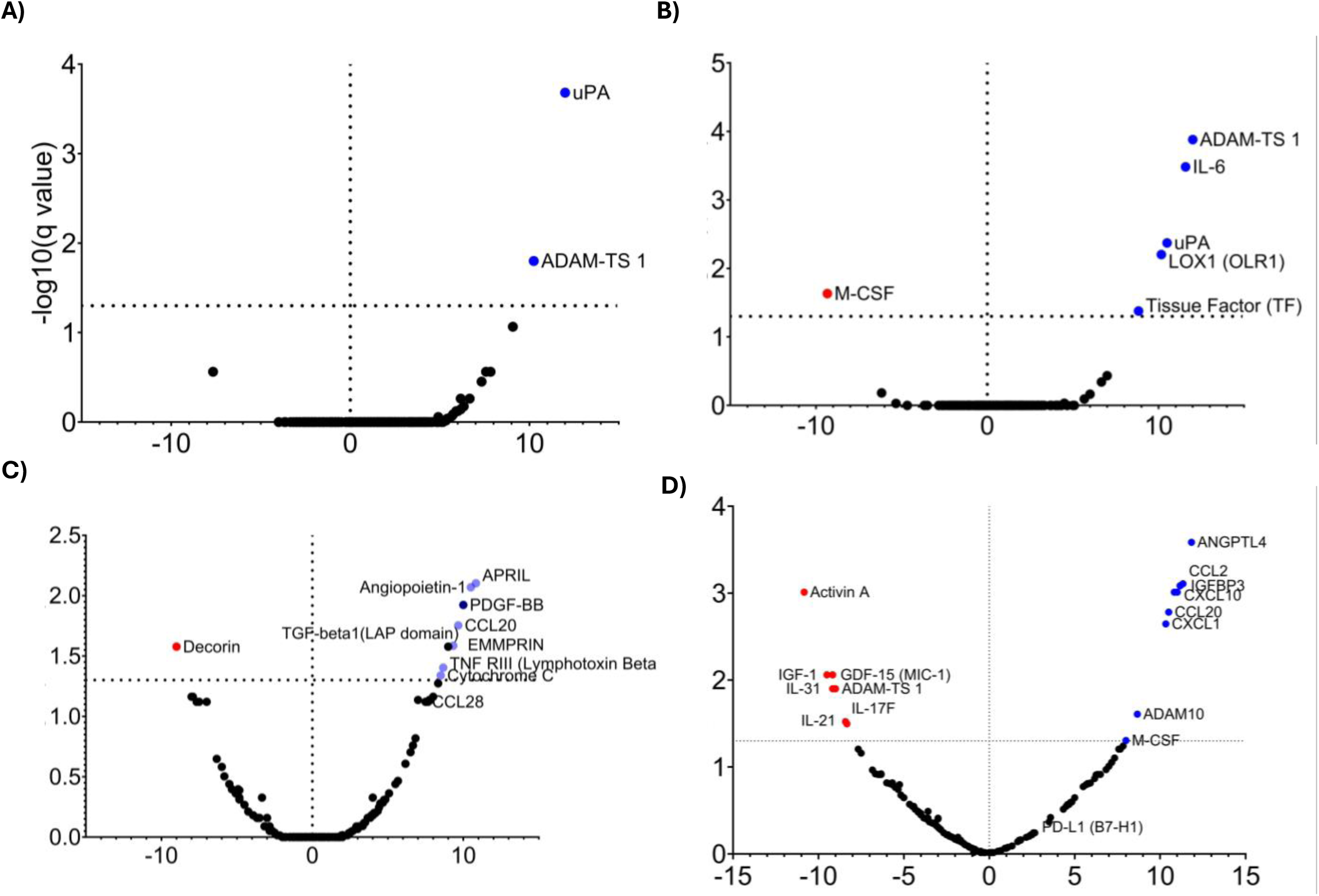
Temporal cytokine profiling reveals dynamic secretome remodeling in MDA-MB-468 cells following Asha therapy: Volcano plots display differential cytokine expression in culture supernatants from Asha-treated versus control cells at **A)** 24 hours **, B)** 48 hours **, C)** 72 hours, and **D)** 96 hours, analyzed using nELISA (270-cytokine panel). The x-axis represents mean rank difference (Asha – control), and the y-axis indicates −log₁₀(FDR) values from Mann–Whitney testing. Cytokines above the horizontal dotted line met the significance threshold (FDR < 0.05). Early time points (24–48 hours) show selective upregulation of ADAMTS1, uPA, and IL-6, whereas later stages (72–96 hours) exhibit broader activation of inflammatory and chemotactic cytokines, including CCL2, CXCL10, CCL20, and CXCL1. The dashed vertical line denotes zero mean rank difference (no change). Each dot represents one cytokine analyzed (n=12 sample / condition / time point).

Notably, IL-6 and ADAMTS1 showed concordant upregulation at both transcript (bulk RNA-seq, 48h) and protein (secreted cytokine) levels, validating the temporal sequence where ER stress-induced transcriptional changes drive sustained secretory remodeling^66^. This progression, from acute stress markers (24-48h) to immune-recruiting chemokines (72-96h), suggests potential for tumor microenvironment remodeling.

### Asha therapy reduced metastases and improved survival in conjunction with anti-PD1 in 4T1

The coordinated induction of ER stress markers and the subsequent emergence of a pro-inflammatory secretory program in vitro, characterized by the release of myeloid and lymphoid chemoattractants (CCL2, CXCL10, and CXCL1), suggested that Asha therapy could remodel the immunosuppressive TNBC TME to sensitize tumors to immune checkpoint blockade. To test this hypothesis, we evaluated Asha therapy in combination with anti-PD1 in the 4T1 orthotopic model.

An initial four-week study evaluated Asha Therapy’s anti-metastatic efficacy in a 4T1 (TNBC) orthotopic syngeneic immunocompetent model. Mice with established tumors were randomized into four groups (n=6/group): control, Asha monotherapy, anti-PD1 monotherapy, and combination therapy. At study completion, lung metastatic nodules were microscopically counted by a trained technician. While Asha monotherapy showed a directional reduction in metastatic burden that did not reach statistical significance, the Asha/anti-PD1 combination achieved significant reduction in lung metastases compared to control (adjusted p=0.037; Figure 4A). These effects occurred without significant changes in primary tumor volume (Supplementary Figure S4A), and the treatment was well-tolerated, with no weight loss observed in any experimental group throughout the study (Supplementary Figure S4B).

**Figure 4.**
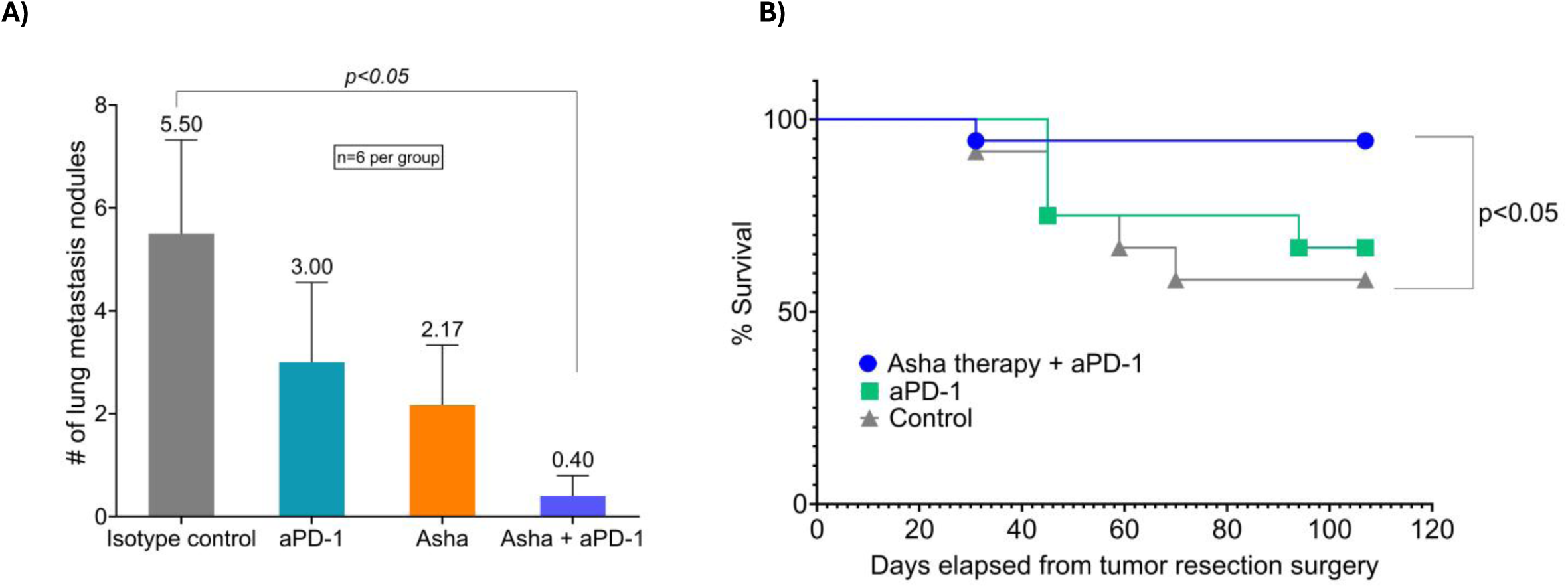
Asha therapy in combination with anti-PD1 significantly reduced metastatic burden, and improved survival in 4T1 TNBC model. **(A)** Lung metastasis quantification in 4T1 tumor-bearing mice following 4-week treatment. Female BALB/c mice with established orthotopic 4T1 tumors were randomized to control (n=6), Asha monotherapy (n=6), anti-PD1 monotherapy (n=6), or combination therapy (n=6). Metastatic nodules were microscopically counted at study completion. Data are shown as individual values with mean ± SEM. Statistical analysis by Kruskal-Wallis test followed by Dunn’s multiple comparisons test **(B)** Kaplan-Meier survival analysis following primary tumor resection. Female BALB/c mice with established 4T1 tumors received 1 week of treatment (control, n=12; anti-PD1 monotherapy, n=12; Asha + anti-PD1 combination, n=18), followed by surgical resection of the primary tumor. Mice were monitored for survival over 107 days post-surgery. Combination therapy significantly reduced mortality risk compared to control (HR = 0.12, 95% CI: 0.02–0.63, p<0.05, log-rank test).

A subsequent study using the 4T1 metastasis model further validated the synergistic effect. Immunocompetent mice with established tumors were randomized into three groups: control (n=12), anti-PD1 monotherapy (n=12), and Asha therapy / anti-PD1 combination (n=18). After one week of treatment, the primary tumor was resected, and the mice were monitored for survival. Anti-PD1 monotherapy showed no significant survival benefit, while the combination therapy reduced death risk by 88% vs control (p<0.05, Hazard ratio = 0.12, 95% CI: 0.02-0.63; Figure 4B). Consistent with our earlier findings, longitudinal monitoring confirmed the safety of the combination regimen, with no evidence of treatment-induced weight loss (Supplementary Figure S4C). These findings demonstrate that Asha therapy converts checkpoint-refractory 4T1 tumors into checkpoint-responsive disease, significantly improving survival when combined with anti-PD1.

### CITE-seq profiling reveals immune reprogramming in the tumor microenvironment

To characterize how the in vitro stress and secretory responses translate to the tumor microenvironment in vivo, we performed CITE-seq profiling of dissociated tumor tissues from control (n=3) and Asha-treated (n=3) mice at 7-days following treatment initiation. Single-cell analysis yielded a median of 4,487 high-quality cells per sample (range: 3,879–5,159). Unsupervised clustering identified 11 distinct populations, which were annotated based on expression of canonical marker gene expression, and confirmed using surface protein markers (Figures 5A-C, Supplementary Figure S5). Subsequent analyses focused on the five most abundant cell types (epithelial/malignant cells, monocytes/macrophages, neutrophils, T cells, and fibroblasts), which collectively represented >90% of profiled cells. Six rare populations (dendritic cells, plasmacytoid dendritic cells, NK cells, endothelial cells, mast cells, and B cells) were catalogued but excluded from differential expression analysis due to insufficient cell numbers. These populations are characteristically rare in the 4T1 TNBC model,^67^ consistent with the poorly immunogenic nature of this tumor.^68^

**Figure 5.**
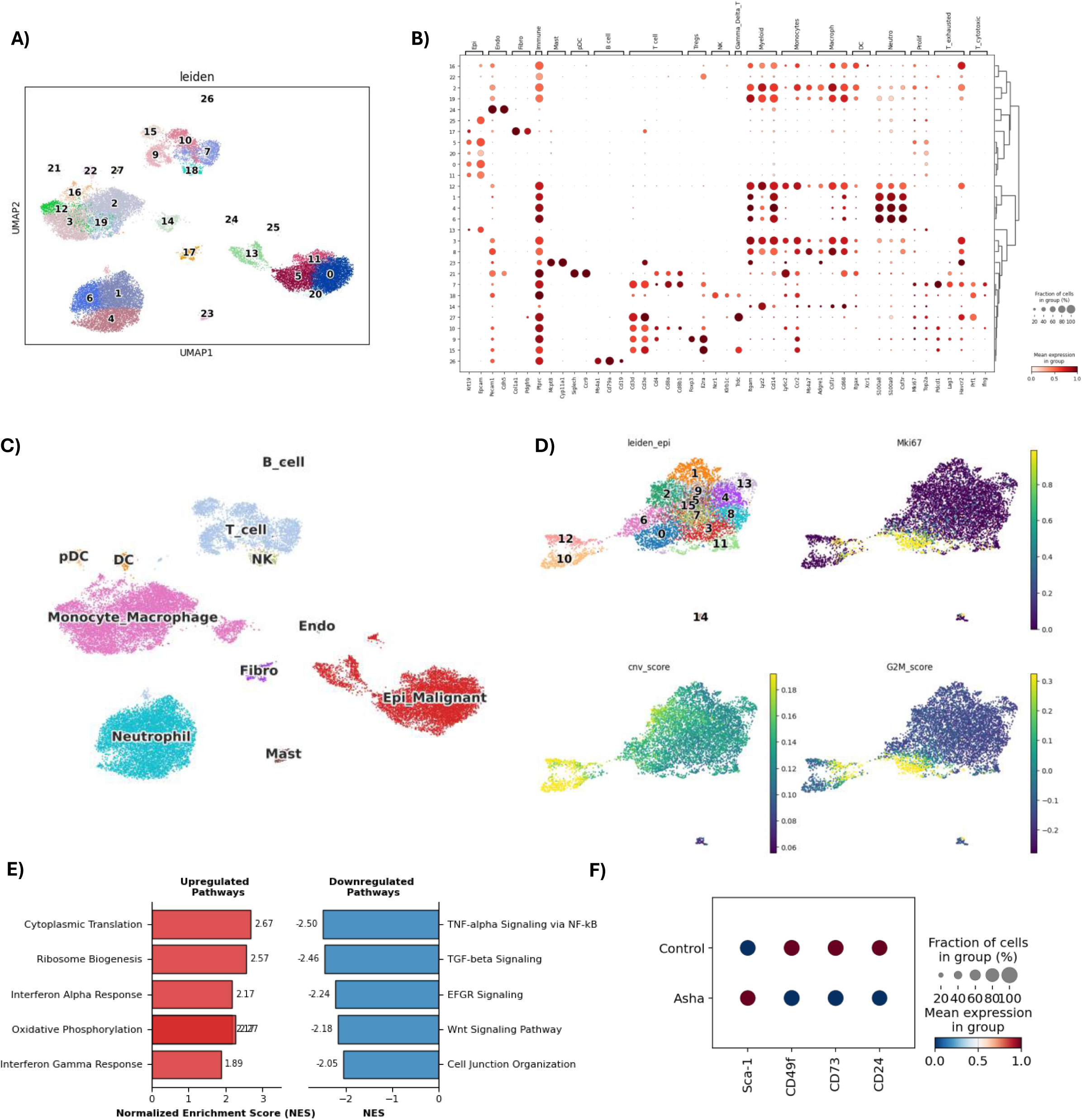
Cellular composition and malignant reprogramming of the 4T1 tumor microenvironment. **(A)** UMAP of 26,446 tumor-infiltrating cells colored by Leiden cluster identity (28 clusters). **(B)** Dot plot showing expression of canonical lineage markers used for cluster identification; dot size indicates the fraction of cells expressing the marker, and color intensity represents mean expression level. **(C)** UMAP colored by annotated cell type (11 populations). Related Leiden clusters were merged based on shared lineage-specific marker expression. **(D)** UMAP plot of sub-clustered epitheilial cells displaying leiden sub-clusters, the proliferation marker Mki67, copy number variation (CNV) scores, and G2M cell cycle scores. Based on elevated Mki67, elevated CNV scores and G2M scores, leiden sub-clusters 0, 10, and 12 were identified as malignant cells. (**E**) Gene Set Enrichment Analysis (GSEA) of malignant cells comparing Asha-treated vs. Control tumors. Bar plot displays representative biological programs significantly enriched (FDR < 0.05) in treated cells, ranked by Normalized Enrichment Score (NES). Upregulated pathways (red) denote a shift toward enhanced biosynthetic and interferon-mediated activity, while downregulated pathways (blue) characterize the loss of oncogenic growth factor responsiveness and structural organization (**F**) Dot plot displaying CITE-seq protein expression for key interferon-response (Sca-1) and immune-evasive/stemness markers (CD49f, CD73, CD24) in the malignant cell compartment. Color indicates the relative mean expression (standardized per marker to highlight shifts), and dot size represents the percentage of cells expressing each marker.

### Asha therapy elicits translational and interferon-driven stress responses in malignant cells

To determine whether the stress responses observed in vitro persist in the complex tumor microenvironment, we analyzed malignant cell transcriptional states within Asha-treated tumors. Epithelial sub-clustering identified approximately 20% of cells as malignant based on high copy number variation, elevated G2M scores, and Mki67 expression (Figure 5D). Restricting differential expression to these high-confidence malignant cells (n=1,394) minimized stromal contamination.

Differential expression revealed upregulation of interferon-induced genes (Ifi27l2a, log₂FC = 2.65) and stress-associated genes (Filip1l, log₂FC = 1.44), alongside the immune checkpoint regulator Cd274/PD-L1 (log₂FC = 1.09). Downregulated genes included stress-responsive transcription factors (Klf2, log₂FC = −1.17) (Supplementary Table S5). GSEA revealed upregulation of biosynthetic programs (cytoplasmic translation, NES = 2.67; ribosome biogenesis, NES = 2.57; oxidative phosphorylation, NES = 2.27; MYC targets, NES = 2.00) alongside interferon signaling (IFN-α, NES = 2.17; IFN-γ, NES = 1.89). Notably, downregulated pathways included TNF-α/NF-κB signaling (NES = −2.50), growth factor pathways (TGF-β, NES = −2.46; EGFR, NES = −2.24; Wnt, NES = −2.18), MAPK cascade (NES = −2.04), and cell junction organization (NES = −2.05) (all FDR < 0.05; Figure 5B; Supplementary Table S6).

Under continuous therapy, malignant cells exhibited interferon-driven and biosynthetic programs without persistent elevation of acute ER stress markers. This pattern suggests cellular adaptation to chronic stress exposure, where the initial ER stress response (documented at 48-96h in vitro) transitions to sustained interferon and metabolic activation. At the protein level, CITE-seq confirmed interferon pathway activation with increased surface expression of the interferon-stimulated gene, Sca-1 (log₂FC = +0.43), alongside reduced expression of the immunosuppression marker CD73 (−0.54) and stemness markers CD49f (−0.93) and CD24 (−0.42), indicating enhanced immunological visibility and reduced immune evasion capacity (Figure 5C; Supplementary Table S7).

### Asha therapy shifts the tumor fibroblast compartment away from a myCAF-like contractile state toward a stress-responsive, immune-recruiting phenotype

Given that our cytokine profiling revealed secretion of factors capable of remodeling the tumor stroma, we examined fibroblast responses to Asha therapy. Differential expression revealed suppression of contractile and matrix associated programs, with downregulation of cytoskeletal genes (Myh11, Mylk, Des), matrix components (Col4a1, Col18a1), and Il6, alongside upregulation of stress-responsive genes (Ankrd1), neutrophil-recruiting chemokines (Cxcl3), and interferon-stimulated genes (Ifit1, Ifi202b) (Supplementary Table S8). GSEA (Figure 6A; Supplementary table S9) showed negative enrichment of epithelial-mesenchymal transition (NES=-2.19) and ECM organization (NES=-2.13), with positive enrichment of ribosomal (NES=2.60), hypoxia (NES=2.21), and interferon responses (NES=2.05-2.00).

**Figure 6.**
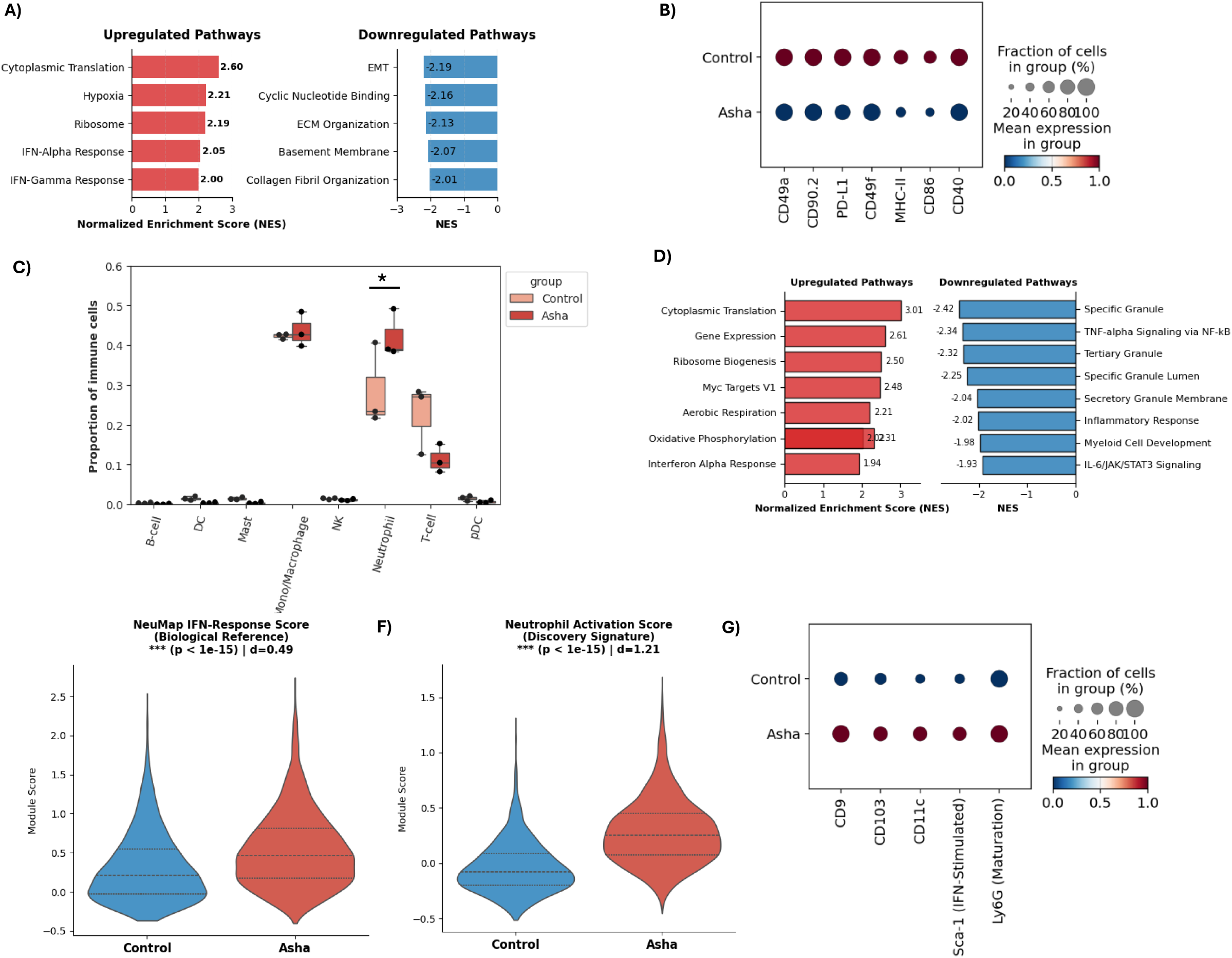
Asha therapy remodels the fibroblasts and drives a specific interferon-responsive neutrophil activation. **(A)** Gene Set Enrichment Analysis (GSEA) of fibroblasts comparing Asha-treated vs. Control tumors. Bar plot displays representative biological programs significantly enriched (FDR < 0.05), ranked by Normalized Enrichment Score (NES). Upregulated pathways (red) signify metabolic reprogramming toward a biosynthetically active (e.g., Cytoplasmic Translation) and interferon-responsive phenotype. Downregulated pathways (blue) demonstrate the suppression of contractile (EMT) and matrix-associated programs (ECM Organization). **(B)** Dot plot displaying CITE-seq protein expression for key structural and immunomodulatory markers in the fibroblasts. Markers include the myofibroblast marker Thy1 (CD90.2), integrin subunits (CD49a, CD61), and the immune-checkpoint ligand PD-L1 (CD274). Color indicates standardized mean expression per marker, and dot size represents the percentage of cells expressing each marker. Asha therapy shifts fibroblasts from a matrix-building state to a stress-responsive, immune-communicating phenotype. **(C)** scCODA compositional analysis showing neutrophils as the sole significantly expanded immune population (log₂FC = 0.81, FDR < 0.05). **(D)** GSEA of neutrophils showing upregulated metabolic/interferon pathways (left, red) and downregulated immunosuppressive/granule pathways (right, blue). NES values indicated on bars; all pathways shown have FDR < 0.05. **(E)** Violin plot of the NeuMap IFN-response module score. Asha therapy drives a specific shift toward the anti-tumor trajectory **(F)** Violin plot of the bespoke Neutrophil Activation Score derived from the top differentially expressed genes in this study. As expected, this discovery signature reveals a profound shift in the treated population compared to control tumors. **(G)** Dotplot showing relative expression of key activation and maturation markers following Asha therapy. Asha therapy drives a robust increase in the surface density of activation markers (CD9, CD103, CD11c) and the interferon-stimulated antigen Sca-1, with a concurrent reduction in the maturation marker Ly6G. Color indicates standardized mean expression per marker, and dot size represents the percentage of cells expressing each marker. For violin plots panels, internal lines represent quartiles and the median; statistical significance was determined by Wilcoxon Rank-Sum test.

CITE-seq showed concordant suppression of contractile surface protein markers, including Thy1/CD90.2 (log₂FC = −0.63), and integrin subunits (CD49a, CD49d, CD61) that mediate matrix adhesion, alongside reduced expression of immune regulatory molecules (MHC-II, CD86, CD40, PD-L1) (Figure 6B; Supplementary Table S10). This stromal reprogramming, contractile suppression with interferon activation, and neutrophil recruitment capacity, may contribute to the anti-metastatic efficacy observed in the 4T1 model.

### Asha therapy increases neutrophil infiltration and reprograms neutrophils toward an interferon-activated state

Neutrophils represent a critical myeloid population capable of both tumor-promoting and anti-tumor functions depending on their activation state. We therefore examined neutrophil responses to Asha therapy. Compositional modeling (scCODA) identified neutrophils as the only immune population significantly increased following Asha therapy (log₂ fold-change = 0.81, FDR < 0.05), while other compartments remained unchanged (Figure 6C). This selective expansion is consistent with recent reports that effective immunotherapies induce systemic proliferation of interferon-stimulated gene (ISG)⁺ anti-tumor neutrophils^23,69^.

Differential gene expression analysis revealed that Asha therapy exposure (Supplementary Table S11) downregulated genes included azurophilic and specific granule components (Ngp, Ltf, Camp, Mmp8, Mmp9, Lcn2) and the immunosuppressive marker Chil3, while upregulating genes included T cell-recruiting chemokines (Cxcl9), and interferon-stimulated genes (Ifit3, Ifi27l2a, Ifi44, Ifi44l). GSEA demonstrated activation of biosynthetic and metabolic programs such as cytoplasmic translation, ribosome biogenesis, oxidative phosphorylation (NES = 2.5–3.0, FDR < 0.001), alongside interferon-alpha response (NES = 1.94, FDR = 0.002), with concurrent suppression of immunosuppressive pathways including TNF-α/NF-κB signaling, and IL-6/JAK/STAT3 signaling (NES = −1.9 to −2.4, FDR < 0.05) (Figure 6D; Supplementary Table S12).

To quantitatively assess the relationship between Asha-induced neutrophils and functionally defined neutrophil states, we calculated module scores for the seven transcriptional hubs mapped in the NeuMap reference atlas^69^. Asha therapy induced a significant enrichment of the IFN-response hub signature (median score: Control = 0.21, Asha = 0.46; p < 10⁻□□, Cohen’s d = 0.49, Figure 6E), a transcriptional state characterized by interferon-stimulated gene expression, metabolic activation, and anti-tumor function. Negligible changes were observed in other hub signatures (Supplementary Figure S6, all Cohen’s d < 0.15). Critically, neutrophils showed no enrichment of immunosuppressive (IS-I, IS-II) or immature hub signatures (p > 0.10), indicating selective activation of the IFN-response trajectory rather than non-specific inflammatory reprogramming. This pattern of interferon activation with suppressed classical inflammatory pathways suggests a shift away from the immunosuppressive, pro-tumorigenic neutrophil states typically observed in untreated tumors. To further characterize this response, we generated a bespoke Neutrophil Activation Score from the top differentially expressed genes in neutrophils exposed to Asha therapy to create a human-projectable signature for clinical cohort validation (Supplementary 1). When applied back to the discovery dataset, this score confirmed the expected shift in the treated population (median score: Control = −0.077, Asha = 0.252; Cohen’s d = 1.21; Figure 6F).

At the protein level, CITE-seq analysis confirmed the transcriptomic findings, showing significant upregulation of activation markers including CD9 (log₂FC = +1.41), CD103 (log₂FC = +0.89), and CD11c (log₂FC = +0.78), as well as the interferon-stimulated surface antigen Ly6A/E (Sca-1; log₂FC = +1.00), with reduced Ly6G (log₂FC = −0.76) (Figure 6G, Supplementary Table S13). The concordance between transcriptomic and proteomic signatures reinforces the interferon-driven activation phenotype.

These findings demonstrate that Asha therapy selectively expands neutrophils and programs them towards a metabolically activated, interferon-responsive state associated with tumor control in immunotherapy contexts. This interferon-driven phenotype contrasts with polymorphonuclear myeloid-derived suppressor cells (PMN-MDSCs), which suppress anti-tumor immunity through arginase-1, ROS, and T cell inhibition, and whose accumulation correlates with poor outcomes in many cancers.

### Asha therapy induces interferon-responsive states in tumor-associated monocytes and macrophages

Macrophages and monocytes are key regulators of tumor immunity and can exhibit either pro-tumor or anti-tumor functions. We examined how Asha therapy affects these populations. Re-clustering identified four mononuclear phagocyte subsets: C1qa⁺Trem2⁺Apoe⁺ lipid-associated macrophages (LAMs, 48.8%), Ly6c2⁺Ccr2⁺ inflammatory monocytes (25.6%), Mrc1⁺Arg1⁺ IL4-responsive macrophages (21.7%), and MHC-II⁺ macrophages (3.8%) (Supplementary Figures S7A-7C). Compositional analysis (scCODA) showed stable subset proportions (Supplementary Figure S7D).

LAMs, the dominant population (48.8%), underwent interferon-driven remodeling with enrichment of type I interferon response (IFN-α, NES = 2.18; IFN-γ, NES = 2.03), antimicrobial defense pathways (NES = 2.29), and neutrophil migration signatures (NES = 1.96), alongside suppressed oxidative phosphorylation (NES = −2.28) and MYC targets (NES = −2.17) (Figure 7A; Supplementary Tables S14-S15). At the protein level, CITE-seq showed increased expression of the interferon-inducible protein Sca-1 (log₂FC = +0.79) but reduced antigen-presentation markers MHC-II (log₂FC = −0.57) and CD86 (−0.39), indicating interferon activation without classical antigen-presenting function (Fig. 7B; Supplementary Table S16).

**Figure 7.**
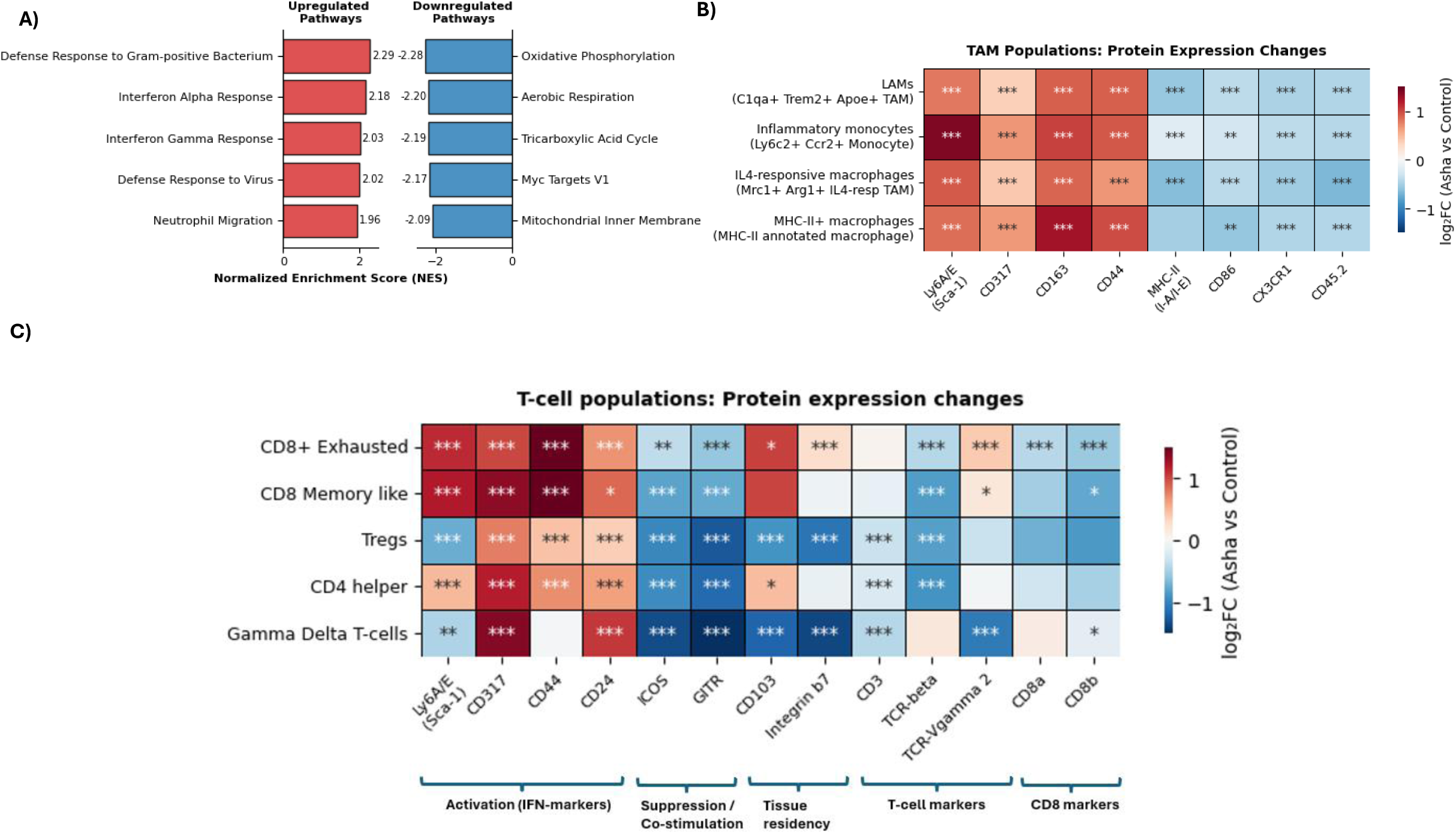
Asha therapy induces inflammatory and metabolic reprogramming in monocytes / macrophages and T-cells. **(A)** Gene Set Enrichment Analysis (GSEA) of the C1qa+ Trem2+ Apoe+ lipid-associated macrophage (LAM) subset following therapy. Bar plots illustrate normalized enrichment scores (NES) for significantly altered pathways (FDR < 0.05). Asha therapy induces interferon-alpha/gamma response and antimicrobial defense pathways alongside neutrophil migration signatures. Conversely, treatment leads to a significant suppression of metabolic and biosynthetic programs, including oxidative phosphorylation, aerobic respiration, and Myc Targets.**(B)** Differential surface protein expression across myeloid subsets. Heatmap shows log₂ (fold-change) (Asha vs Control) for TAM subsets: C1qa⁺Trem2⁺Apoe⁺ lipid-associated macrophages (LAMs), Ly6c2⁺Ccr2⁺ inflammatory monocytes, Mrc1⁺Arg1⁺ IL4-responsive macrophages, and MHC-II⁺ macrophages. Red indicates upregulation, blue indicates downregulation. Note the universal upregulation of the interferon-responsive marker Ly6A/E (Sca-1) and CD317, alongside the downregulation of homeostatic antigen-presentation (MHC-II, CD86) **(C)** Differential surface protein expression across T-cell subsetsHeatmap shows log₂ (fold-change) (Asha vs Control) for T cell subsets: exhausted CD8⁺ T cells, regulatory T cells (Tregs), CD4⁺ helper T cells, CD8⁺ memory-like T cells, and gamma delta T cells. Treatment induced universal upregulation of the interferon-inducible marker CD317 and broad downregulation of co-stimulatory molecules (GITR, ICOS, CD52) and TCR-beta across all populations. Sca-1 (Ly6A/E) increased selectively in CD8⁺ and CD4⁺ subsets but decreased in regulatory T cells. Exhausted CD8⁺ cells showed increased activation markers (CD3, CD44) with reduced CD8a/b expression, regulatory T cells lost canonical suppressive markers (GITR, ICOS, CD103), and Gamma Delta T cells exhibited dramatic reductions in tissue residency markers (integrin β7, CD103) and TCR Vgamma2. Significance was determined via Wilcoxon rank-sum test with Benjamini-Hochberg correction in heatmaps, ***, **, and * correspond to p < 0.001, p < 0.01, and p < 0.05, respectively.

IL4-responsive macrophages showed similar interferon-associated remodeling (increased Sca-1, +0.91; reduced MHC-II, −0.64; CX3CR1, −0.53) with enrichment of alarmin-driven pathways (S100a8, S100a9; NES = 2.39) (Figure 7B, Supplementary Tables S14-S16). Inflammatory monocytes and MHC-II⁺ macrophages exhibited protein-level interferon signatures (increased Sca-1, CD317; reduced CX3CR1) with minimal transcriptional changes (Figure 7B, Supplementary Tables S14-S16). Together with the observed neutrophil expansion (Fig. 6A), these findings indicate interferon-mediated coordination of innate immune compartments within the tumor microenvironment.

### Asha therapy drives systemic biosynthesis and metabolic reprogramming of the T cell compartment

Re-clustering of the T-cell compartment identified five transcriptionally distinct subsets: exhausted CD8⁺ (29.1%), regulatory T cells (26.3%), CD4⁺ helper (18.3%), CD8⁺ memory-like (13.0%), and γδ T cells (13.4%) (Supplementary Figure S8A-C). Compositional analysis (scCODA) showed stable subset proportions (Supplementary Figure S8D).

All αβ T-cell populations showed robust biosynthetic activation with strong enrichment of translational pathways (cytoplasmic translation and ribosome biogenesis, NES = 2.1–3.2, FDR < 0.001). Despite shared metabolic activation, functional responses diverged across subsets. Exhausted CD8⁺ T cells uniquely coupled biosynthetic priming with inflammatory effector activation, showing enrichment of TNF-α/NF-κB signaling (NES = 2.33) and upregulation of Ifng (log₂FC = +1.82), alongside increased surface activation markers (CD3, CD44, Sca-1) (Fig. 7C; Supplementary Table S17-19). In contrast, CD4⁺ helper and CD8⁺ memory-like cells exhibited strong ribosomal enrichment (NES = 2.7–3.2) but suppressed TCR signaling (NES = −2.2 to − 2.3) and reduced co-stimulatory markers (GITR, ICOS, TCRβ), consistent with metabolic priming without effector engagement (Fig. 7C, Supplementary Tables S17-S19).

Regulatory T cells showed biosynthetic and inflammatory activation (TNF-α/NF-κB, NES = 2.05; IFN-α response, NES = 2.00) but lost suppressive surface markers GITR (log₂FC = −1.27, padj < 10⁻¹□□), ICOS (−0.97), and CD103 (−0.90), indicating reduced immunosuppressive capacity (Fig. 7C, Supplementary Tables S17-S19). γδ T cells displayed interferon-driven stress remodeling (IFN-α response, NES = 2.44) without ribosomal activation, with dramatic reductions in tissue residency markers (integrin β7, −1.37; CD103, −1.20; TCR Vγ2, −1.06) and co-stimulatory molecules (GITR, −1.68; ICOS, −1.35).

These findings demonstrate selective CD8⁺ effector activation and reduced Treg suppression within a broadly reprogrammed T-cell compartment, with CD4⁺ and CD8⁺ memory populations primed but requiring additional signals, consistent with the checkpoint blockade synergy observed in vivo.

### Projection of the neutrophil ISG program onto human TNBC tumors

To test whether the neutrophil program we observed in mice also appears relevant in human breast cancer, we projected our 19-gene neutrophil ISG signature onto the METABRIC TNBC cohort (n = 338) and SCANB cohort (n=874). In METABRIC and SCAN-B, high NSG score was associated with significantly improved overall survival by Kaplan–Meier analysis (Figure 8A-8B, log-rank p= 0.036 and and p=0.048). In multivariable Cox models adjusting for nodal status, T stage, tumor size, and stratifying by age, both cohorts showed a consistent protective effect (METABRIC: HR=0.65, 95% CI 0.41–1.03, p=0.063; SCAN-B: HR=0.79, 95% CI 0.60–1.03, p=0.080; Figure 8C), although individual studies were modestly underpowered for fully definitive adjusted significance. When the two cohorts were analyzed jointly in a pooled Cox model with study as a stratification factor and the same covariate structure, the association reached statistical significance (Figure 8C, pooled HR=0.75, 95% CI 0.59–0.94, p=0.01), indicating a robust, reproducible survival benefit of high NSG score across datasets.

**Figure 8.**
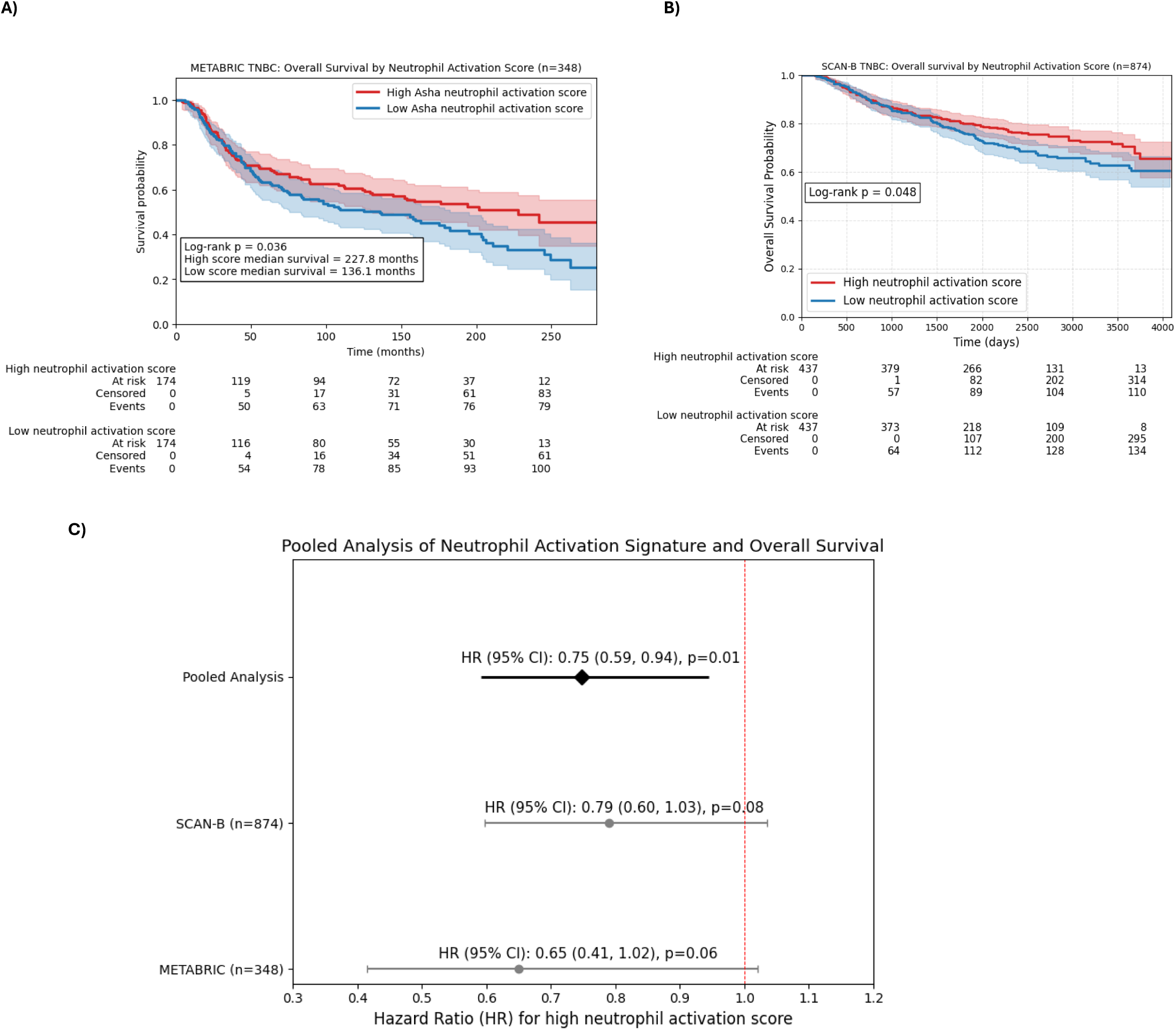
Clinical validation of the Neutrophil Activation Signature in human TNBC studies. **(A)** Kaplan-Meier survival analysis of the METABRIC cohort. Triple-negative breast cancer (TNBC) patients (n=348) were stratified by the median expression of the Neutrophil Activation Signature. High signature expression is significantly associated with improved overall survival (Log-rank p=0.036). **(B)** Kaplan-Meier survival analysis of the SCAN-B cohort. Independent validation in the SCAN-B TNBC population (n=874) confirms that a high Neutrophil Activation Score is a significant predictor of favorable patient outcomes (Log-rank p=0.048). **(C)** Multivariable adjusted pooled analysis of independent prognostic value. Forest plot displays the Hazard Ratios (HR) for the individual cohorts and the pooled analysis, derived from multivariable Cox proportional hazards models adjusting for nodal status, T stage, tumor size, and age. While individual cohorts showed a consistent protective trend, the pooled analysis (total n = 1,222) demonstrates that the Neutrophil Activation Signature is a significant independent predictor of favorable survival (pooled HR = 0.75; 95% CI: 0.59, 0.94; p = 0.01). Pooled HR was calculated using a fixed-effect model with study as a stratification factor. In Kaplan-Meier curves, solid lines represent the survival probability, and shaded areas indicate the 95% confidence intervals.

## Discussion

Using multi-omics analysis across in vitro models and in vivo tumor microenvironments, we demonstrate that low-intensity EMF-based Asha therapy is associated with coordinated tumor and stromal reprogramming that correlates with reduced metastatic progression, and enhanced responsiveness to the immune checkpoint blockade.

### Asha therapy shifts malignant cells to an immunogenic stress state

A key tumor-intrinsic response to Asha therapy is the induction of adaptive ER stress. Unlike traditional cytotoxic agents, our data suggest Asha therapy acts as a reprogramming modality; it activates canonical UPR pathways without inducing acute cytotoxicity. This stress response is not static; it matures from an initial disruption of protein folding accompanied by secretion of immune-recruiting factors. Within 48–96 hours, tumor cells fundamentally rewire their internal signaling, characterized by a reciprocal downregulation of anabolic mTOR effectors and a massive upregulation of the stress-adaptive cAMP-PKA pathway. The dramatic induction of ITPKA suggests that calcium signaling disruption may serve as the biophysical bridge between electromagnetic exposure and this cellular stress state.

This stress program evolves under chronic *in vivo* exposure. While acute *in vitro* responses are dominated by NF-kb activation, malignant cells after seven days of continuous treatment exhibit a transition toward stable, interferon-driven states with suppressed NF-κB signaling. The temporal dynamics of this NF-κB response, activation during acute stress followed by suppression under chronic exposure, suggest adaptation from pro-inflammatory to interferon-dominant signaling. Crucially, this is not a state of passive tolerance, but an active metabolic and biosynthetic reprogramming. These cells maintain high metabolic activity while simultaneously displaying enhanced immunological visibility through increased MHC class II expression and the loss of immunosuppressive markers like CD73 and the stemness-associated factors CD49f and CD24. The persistence of these shifts suggests that Asha therapy maintains the tumor in a chronically altered state, effectively trading its baseline homeostatic profile for an immunogenic phenotype that is more susceptible to immune surveillance

### Selective Neutrophil Reprogramming as a key feature of the Asha Therapy response

Among all immune populations in the TME, neutrophils exhibited the most profound response to Asha therapy, characterized by both a significant compositional expansion and a phenotypic shift. This reprogramming closely mirrors recently described interferon-activated anti-tumor states^23,69^. By mapping our data against established neutrophil atlases, we confirmed a selective activation of these anti-tumor programs rather than a nonspecific inflammatory response. These findings identify neutrophil reprogramming as a prominent feature of the Asha therapy-induced immune response. The clinical relevance of this neutrophil program is supported by its association with improved outcomes in human TNBC. A 19-gene activation signature, derived directly from Asha-treated murine neutrophils, identified patients with significantly improved overall survival across two independent cohorts (METABRIC, n=338; SCAN-B, n=874). This cross-species conservation, where a murine treatment-induced signature successfully stratifies human outcomes, supports the biological relevance of the Asha-induced neutrophil state. These results suggest that targeting or monitoring this specific neutrophil program could offer a new therapeutic or biomarker-driven roadmap for managing aggressive breast cancers.

### Coordinated Stromal and Myeloid Reprogramming

Asha therapy reshapes the TME **t**hrough both compositional and functional changes in myeloid populations. Neutrophils expanded significantly, while macrophages and monocytes underwent transcriptional remodeling without changes in relative abundance. This coordinated reprogramming of the TME is characterized by an interferon-mediated shift across the stromal and myeloid compartments. Cancer-associated fibroblasts transitioned away from contractile, matrix-depositing programs, which typically facilitate metastasis, toward stress-responsive phenotypes with enhanced neutrophil-recruitment capacity. Simultaneously, tumor-associated macrophages adopted an innate inflammatory profile characterized by interferon-inducible stress responses despite reduced antigen-presentation markers (MHC-II, CD86).

This shift in the TME creates a "priming" effect that lowers the activation threshold for adaptive immunity. We observed widespread biosynthetic and metabolic activation across the T-cell compartment, placing CD4+ and memory-like CD8+ T-cells in a "ready" state despite a lack of immediate effector engagement. Furthermore, a reduction in suppressive markers on regulatory T-cells suggests a localized weakening of immune brakes. This coordinated priming provides the mechanistic rationale for the significantly reduced lung metastasis, and 88% reduction in death risk (HR=0.12) observed when Asha therapy is combined with anti-PD1. By transforming a contractile, immunosuppressive stroma into an interferon-rich, metabolically active environment, Asha therapy appears to enhance responsiveness to checkpoint blockade specifically in metastatic disease. Notably, these effects occurred without significant primary tumor growth inhibition, suggesting that Asha-induced immune activation may be differentially effective in the metastatic niche. Whether this reflects differences in tumor architecture, immune composition, or other microenvironmental factors between primary and metastatic sites remains to be determined. When combined with anti-PD1, Asha therapy significantly reduced metastatic burden and mortality, indicating that TME reprogramming can enhance checkpoint efficacy even in settings where primary tumor control is limited.

### Asha therapy represents a non-cytotoxic paradigm for EMF-based oncology

The mechanism of Asha therapy is fundamentally distinct from existing physical tumor interventions. While Tumor Treating Fields (TTFields) achieve anti-tumor effects through mitotic disruption at higher frequencies^70^, and oncomagnetic therapies focus on acute mitochondrial ROS production^30^, Asha’s low-intensity fields are associated with a stable, adaptive stress that reprograms rather than destroys cells. This distinction is clinically significant; by avoiding direct cytotoxicity, Asha Therapy potentially bypasses the dose-limiting toxicities and resistance challenges common to traditional modalities. The temporal adaptation we observed suggests that Asha therapy is associated with maintaining the tumor microenvironment in a continuously altered state, offering a controllable and reversible intervention that may affect even therapy-resistant subpopulations.

This reprogramming capacity has direct translational implications for overcoming primary checkpoint resistance in triple-negative breast cancer (TNBC). The conversion of checkpoint-refractory tumors into responsive disease through electromagnetic field priming addresses a critical unmet need for patients who derive limited benefit from current checkpoint inhibitors. Our findings suggest that Asha therapy may be most effective in tumors with baseline infiltration, where it can activate the resident immune compartment. Furthermore, the neutrophil-centric reprogramming identified here offers a potential roadmap for clinical monitoring: interferon-stimulated gene signatures in peripheral blood or biopsies could serve as potential pharmacodynamic biomarkers to guide therapy. While continuous administration in mice was well-tolerated without overt toxicity, formal dose-ranging and toxicity studies in normal tissues will be essential for clinical translation. Taken together these findings suggest Asha therapy as a candidate non-pharmacologic strategy to sensitize aggressive tumors to immunotherapy.

### Limitations and future directions

While our findings establish a mechanistic framework for Asha therapy, several questions remain. The precise biophysical link between low-intensity EMF exposure and ER calcium disruption requires further validation through direct calcium imaging and genetic perturbation of identified hubs like ITPKA. Similarly, the temporal transition from acute NF-kb activation to a sustained interferon state likely involves intermediate regulatory steps that, if identified, could be exploited to optimize dosing protocols.

Our efficacy studies were restricted to the 4T1 murine model and combination with anti-PD1 blockade. While the cross-species conservation of the ER stress and neutrophil signatures supports translational relevance, future work is needed to evaluate Asha therapy across the full spectrum of human TNBC heterogeneity, particularly in immune-excluded "cold" tumors. Exploring synergy with standard-of-care chemotherapy or alternative immunotherapies, such as CTLA-4 blockade, remains a critical translational priority. Finally, systematic optimization of field parameters (including frequency, intensity, and duty cycle) will be essential to maximize the depth of microenvironment reprogramming in human patients. Addressing these areas will be vital to moving EMF-based immune priming from a preclinical discovery toward a viable clinical intervention for aggressive cancers.

## Conclusions

This work establishes a mechanistic framework for using Asha therapy for modulating TME in TNBC. Asha therapy’s low-intensity EMF induced adaptive ER stress in tumor cells, initiating a cascade of microenvironment reprogramming characterized by selective neutrophil expansion and interferon-driven activation, fibroblast phenotypic switching, and T cell metabolic priming. Combined with checkpoint blockade, these changes achieve substantial therapeutic benefit, providing rationale for clinical development of Asha therapy as an immune priming strategy in triple-negative breast cancer.

The temporal evolution from acute stress to TME reprogramming, and the neutrophil-centric coordination of anti-tumor immunity represent novel insights into how EMF-based modalities can reprogram tumor microenvironments. These findings position Asha therapy as a mechanistically grounded immunomodulatory strategy with standalone anti-tumor activity and the capacity to enhance checkpoint responsiveness in TNBC and other solid tumors.

## References

1. Sharma, P. & Allison, J. P. Immune Checkpoint Targeting in Cancer Therapy: Toward Combination Strategies with Curative Potential. Cell 161, 205–214 (2015).

2. Robert, C. et al. Ipilimumab plus Dacarbazine for Previously Untreated Metastatic Melanoma. N. Engl. J. Med. 364, 2517–2526 (2011).

3. Hamid, O. et al. Safety and Tumor Responses with Lambrolizumab (Anti–PD-1) in Melanoma. N. Engl. J. Med. 369, 134–144 (2013).

4. Reck, M. et al. Pembrolizumab versus Chemotherapy for PD-L1–Positive Non–Small-Cell Lung Cancer. N. Engl. J. Med. 375, 1823–1833 (2016).

5. Le, D. T. et al. Mismatch repair deficiency predicts response of solid tumors to PD-1 blockade. Science 357, 409–413 (2017).

6. Schmid, P. et al. Atezolizumab and Nab-Paclitaxel in Advanced Triple-Negative Breast Cancer. N. Engl. J. Med. 379, 2108–2121 (2018).

7. Burtness, B. et al. Pembrolizumab alone or with chemotherapy versus cetuximab with chemotherapy for recurrent or metastatic squamous cell carcinoma of the head and neck (KEYNOTE-048): a randomised, open-label, phase 3 study. Lancet Lond. Engl. 394, 1915–1928 (2019).

8. Rosenberg, J. E. et al. Atezolizumab in patients with locally advanced and metastatic urothelial carcinoma who have progressed following treatment with platinum-based chemotherapy: a single-arm, multicentre, phase 2 trial. The Lancet 387, 1909–1920 (2016).

9. Schmid, P. et al. Pembrolizumab for Early Triple-Negative Breast Cancer. N. Engl. J. Med. 382, 810–821 (2020).

10. Prasad, V., Haslam, A. & Olivier, T. Updated estimates of eligibility and response: Immune checkpoint inhibitors. J. Clin. Oncol. 42, e14613–e14613 (2024).

11. Tan, S., Li, D. & Zhu, X. Cancer immunotherapy: Pros, cons and beyond. Biomed. Pharmacother. 124, 109821 (2020).

12. Larkin, J. et al. Five-Year Survival with Combined Nivolumab and Ipilimumab in Advanced Melanoma. N. Engl. J. Med. 381, 1535–1546 (2019).

13. Wolchok, J. D. et al. Overall Survival with Combined Nivolumab and Ipilimumab in Advanced Melanoma. N. Engl. J. Med. 377, 1345–1356 (2017).

14. Cortes, J. et al. Pembrolizumab plus Chemotherapy in Advanced Triple-Negative Breast Cancer. N. Engl. J. Med. 387, 217–226 (2022).

15. Lin, N. U. et al. Sites of distant recurrence and clinical outcomes in patients with metastatic triple-negative breast cancer: high incidence of central nervous system metastases. Cancer 113, 2638–2645 (2008).

16. Dent, R. et al. Triple-negative breast cancer: clinical features and patterns of recurrence. Clin. Cancer Res. Off. J. Am. Assoc. Cancer Res. 13, 4429–4434 (2007).

17. Binnewies, M. et al. Understanding the tumor immune microenvironment (TIME) for effective therapy. Nat. Med. 24, 541–550 (2018).

18. Mellman, I., Chen, D. S., Powles, T. & Turley, S. J. The cancer-immunity cycle: Indication, genotype, and immunotype. Immunity 56, 2188–2205 (2023).

19. Tumeh, P. C. et al. PD-1 blockade induces responses by inhibiting adaptive immune resistance. Nature 515, 568–571 (2014).

20. Twyman-Saint Victor, C., et al. Radiation and dual checkpoint blockade activate non-redundant immune mechanisms in cancer. Nature 520, 373–377 (2015).

21. Ribas, A. et al. Oncolytic Virotherapy Promotes Intratumoral T Cell Infiltration and Improves Anti-PD-1 Immunotherapy. Cell 170, 1109–1119.e10 (2017).

22. Sharma, M. et al. Bempegaldesleukin selectively depletes intratumoral Tregs and potentiates T cell-mediated cancer therapy. Nat. Commun. 11, 661 (2020).

23. Gungabeesoon, J. et al. A neutrophil response linked to tumor control in immunotherapy. Cell 186, 1448–1464.e20 (2023).

24. Hirschhorn, D. et al. T cell immunotherapies engage neutrophils to eliminate tumor antigen escape variants. Cell 186, 1432–1447.e17 (2023).

25. Stupp, R. et al. Effect of Tumor-Treating Fields Plus Maintenance Temozolomide vs Maintenance Temozolomide Alone on Survival in Patients With Glioblastoma: A Randomized Clinical Trial. JAMA 318, 2306–2316 (2017).

26. Ceresoli, G. L. et al. Tumour Treating Fields in combination with pemetrexed and cisplatin or carboplatin as first-line treatment for unresectable malignant pleural mesothelioma (STELLAR): a multicentre, single-arm phase 2 trial. Lancet Oncol. 20, 1702–1709 (2019).

27. Leal, T. et al. Tumor Treating Fields therapy with standard systemic therapy versus standard systemic therapy alone in metastatic non-small-cell lung cancer following progression on or after platinum-based therapy (LUNAR): a randomised, open-label, pivotal phase 3 study. Lancet Oncol. 24, 1002–1017 (2023).

28. Voloshin, T. et al. Tumor-treating fields (TTFields) induce immunogenic cell death resulting in enhanced antitumor efficacy when combined with anti-PD-1 therapy. Cancer Immunol. Immunother. CII 69, 1191–1204 (2020).

29. Lin, W. et al. Tumor treating fields enhance anti-PD therapy by improving CCL2/8 and CXCL9/CXCL10 expression through inducing immunogenic cell death in NSCLC models. BMC Cancer 25, 489 (2025).

30. Hambarde, S., Manalo, J. M., Baskin, D. S., Sharpe, M. A. & Helekar, S. A. Spinning magnetic field patterns that cause oncolysis by oxidative stress in glioma cells. Sci. Rep. 13, 19264 (2023).

31. Helekar, S., Hambarde, S., Baskin, D. & Sharpe, M. EXTH-13. POTENT ANTICANCER EFFECTS OF A NEW WEARABLE NONINVASIVE ONCOMAGNETIC DEVICE: CELLULAR MECHANISMS OF ACTION. Neuro-Oncol. 22, ii89 (2020).

32. Charan, M. et al. Induced electric fields inhibit breast cancer growth and metastasis by modulating the immune tumor microenvironment. 2024.04.14.589256 Preprint at 10.1101/2024.04.14.589256 (2024).

33. Kirimlioglu, E. et al. Short and long-term 2100 MHz radiofrequency radiation causes endoplasmic reticulum stress in rat testis. Histochem. Cell Biol. 162, 311–321 (2024).

34. Blank, M. & Goodman, R. Electromagnetic fields stress living cells. Pathophysiology 16, 71–78 (2009).

35. Furumoto, Y. et al. Activation of Endoplasmic Reticulum Stress Response by Applying of Nanosecond Pulsed Electric Fields for Medical Application. in 2018 IEEE International Power Modulator and High Voltage Conference (IPMHVC) 456–460 (2018). doi:10.1109/IPMHVC.2018.8936739.

36. Zhang, K. & Kaufman, R. J. From endoplasmic-reticulum stress to the inflammatory response. Nature 454, 455–462 (2008).

37. Rufo, N., Garg, A. D. & Agostinis, P. The Unfolded Protein Response in Immunogenic Cell Death and Cancer Immunotherapy. Trends Cancer 3, 643–658 (2017).

38. Chen, S., Zhou, Y., Chen, Y. & Gu, J. fastp: an ultra-fast all-in-one FASTQ preprocessor. Bioinformatics 34, i884–i890 (2018).

39. Dobin, A. et al. STAR: ultrafast universal RNA-seq aligner. Bioinformatics 29, 15–21 (2013).

40. Love, M. I., Huber, W. & Anders, S. Moderated estimation of fold change and dispersion for RNA-seq data with DESeq2. Genome Biol. 15, 550 (2014).

41. Muzellec, B., Teleńczuk, M., Cabeli, V. & Andreux, M. PyDESeq2: a python package for bulk RNA-seq differential expression analysis. Bioinforma. Oxf. Engl. 39, btad547 (2023).

42. Subramanian, A. et al. Gene set enrichment analysis: A knowledge-based approach for interpreting genome-wide expression profiles. Proc. Natl. Acad. Sci. 102, 15545–15550 (2005).

43. Liberzon, A. et al. The Molecular Signatures Database (MSigDB) hallmark gene set collection. Cell Syst. 1, 417–425 (2015).

44. Dagher, M. et al. nELISA: a high-throughput, high-plex platform enables quantitative profiling of the inflammatory secretome. Nat. Methods 1–11 (2025) doi:10.1038/s41592-025-02861-6.

45. Langley, S. R. & Mayr, M. Comparative analysis of statistical methods used for detecting differential expression in label-free mass spectrometry proteomics. J. Proteomics 129, 83–92 (2015).

46. Zheng, H., Siddharth, S., Parida, S., Wu, X. & Sharma, D. Tumor Microenvironment: Key Players in Triple Negative Breast Cancer Immunomodulation. Cancers 13, 3357 (2021).

47. Fleming, S. J. et al. Unsupervised removal of systematic background noise from droplet-based single-cell experiments using CellBender. Nat. Methods 20, 1323–1335 (2023).

48. Wolf, F. A., Angerer, P. & Theis, F. J. SCANPY: large-scale single-cell gene expression data analysis. Genome Biol. 19, 15 (2018).

49. Wolock, S. L., Lopez, R. & Klein, A. M. Scrublet: Computational Identification of Cell Doublets in Single-Cell Transcriptomic Data. Cell Syst. 8, 281–291.e9 (2019).

50. Gayoso, A. et al. Joint probabilistic modeling of single-cell multi-omic data with totalVI. Nat. Methods 18, 272–282 (2021).

51. McInnes, L., Healy, J., Saul, N. & Großberger, L. UMAP: Uniform Manifold Approximation and Projection. J. Open Source Softw. 3, 861 (2018).

52. Traag, V. A., Waltman, L. & van Eck, N. J. From Louvain to Leiden: guaranteeing well-connected communities. Sci. Rep. 9, 5233 (2019).

53. Büttner, M., Ostner, J., Müller, C. L., Theis, F. J. & Schubert, B. scCODA is a Bayesian model for compositional single-cell data analysis. Nat. Commun. 12, 6876 (2021).

54. Squair, J. W. et al. Confronting false discoveries in single-cell differential expression. Nat. Commun. 12, 5692 (2021).

55. Zimmerman, K. D., Espeland, M. A. & Langefeld, C. D. A practical solution to pseudoreplication bias in single-cell studies. Nat. Commun. 12, 738 (2021).

56. Lin, M.-H. et al. Benchmarking differential expression, imputation and quantification methods for proteomics data. Brief. Bioinform. 23, bbac138 (2022).

57. Patel, A. P. et al. Single-cell RNA-seq highlights intratumoral heterogeneity in primary glioblastoma. Science 344, 1396–1401 (2014).

58. Lehmann, B. D. et al. Multi-omics analysis identifies therapeutic vulnerabilities in triple-negative breast cancer subtypes. Nat. Commun. 12, 6276 (2021).

59. Veglia, F. et al. Analysis of classical neutrophils and polymorphonuclear myeloid-derived suppressor cells in cancer patients and tumor-bearing mice. J. Exp. Med. 218, e20201803 (2021).

60. Lee, A. S. The ER chaperone and signaling regulator GRP78/BiP as a monitor of endoplasmic reticulum stress. Methods San Diego Calif 35, 373–381 (2005).

61. Walter, P. & Ron, D. The unfolded protein response: from stress pathway to homeostatic regulation. Science 334, 1081–1086 (2011).

62. Puthalakath, H. et al. ER Stress Triggers Apoptosis by Activating BH3-Only Protein Bim. Cell 129, 1337–1349 (2007).

63. Deshmane, S. L., Kremlev, S., Amini, S. & Sawaya, B. E. Monocyte chemoattractant protein-1 (MCP-1): an overview. J. Interferon Cytokine Res. Off. J. Int. Soc. Interferon Cytokine Res. 29, 313–326 (2009).

64. Luster, A. D. Chemokines — Chemotactic Cytokines That Mediate Inflammation. N. Engl. J. Med. 338, 436–445 (1998).

65. De Filippo, K. et al. Mast cell and macrophage chemokines CXCL1/CXCL2 control the early stage of neutrophil recruitment during tissue inflammation. Blood 121, 4930–4937 (2013).

66. Freitas, V. M. et al. Decreased expression of ADAMTS-1 in human breast tumors stimulates migration and invasion. Mol. Cancer 12, 2 (2013).

67. Ma, Q. et al. CXCL13 expression in mouse 4T1 breast cancer microenvironment elicits antitumor immune response by regulating immune cell infiltration. Precis. Clin. Med. 4, 155–167 (2021).

68. Demaria, S. et al. Immune-mediated inhibition of metastases after treatment with local radiation and CTLA-4 blockade in a mouse model of breast cancer. Clin. Cancer Res. Off. J. Am. Assoc. Cancer Res. 11, 728–734 (2005).

69. Cerezo-Wallis, D. et al. Architecture of the neutrophil compartment. Nature 1–10 (2025) doi:10.1038/s41586-025-09807-0.

70. Kirson, E. D. et al. Disruption of Cancer Cell Replication by Alternating Electric Fields. Cancer Res. 64, 3288–3295 (2004).

