## Supplementary figures for "Magnetic field-induced ER stress reprograms the tumor microenvironment to improve triple-negative breast cancer survival"

### Slide 1
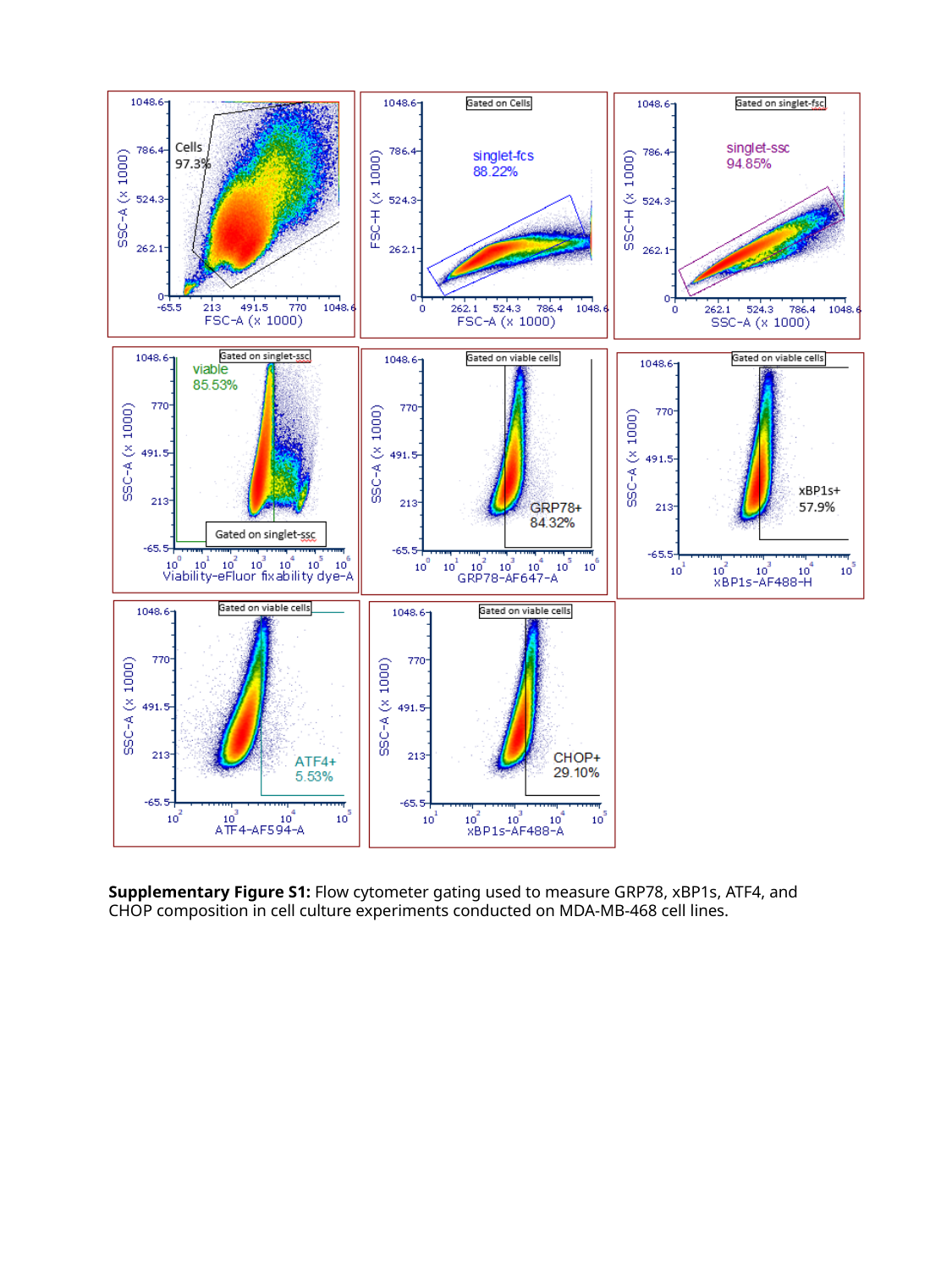

Supplementary Figure S1: Flow cytometer gating used to measure GRP78, xBP1s, ATF4, and CHOP composition in cell culture experiments conducted on MDA-MB-468 cell lines.

### Slide 2
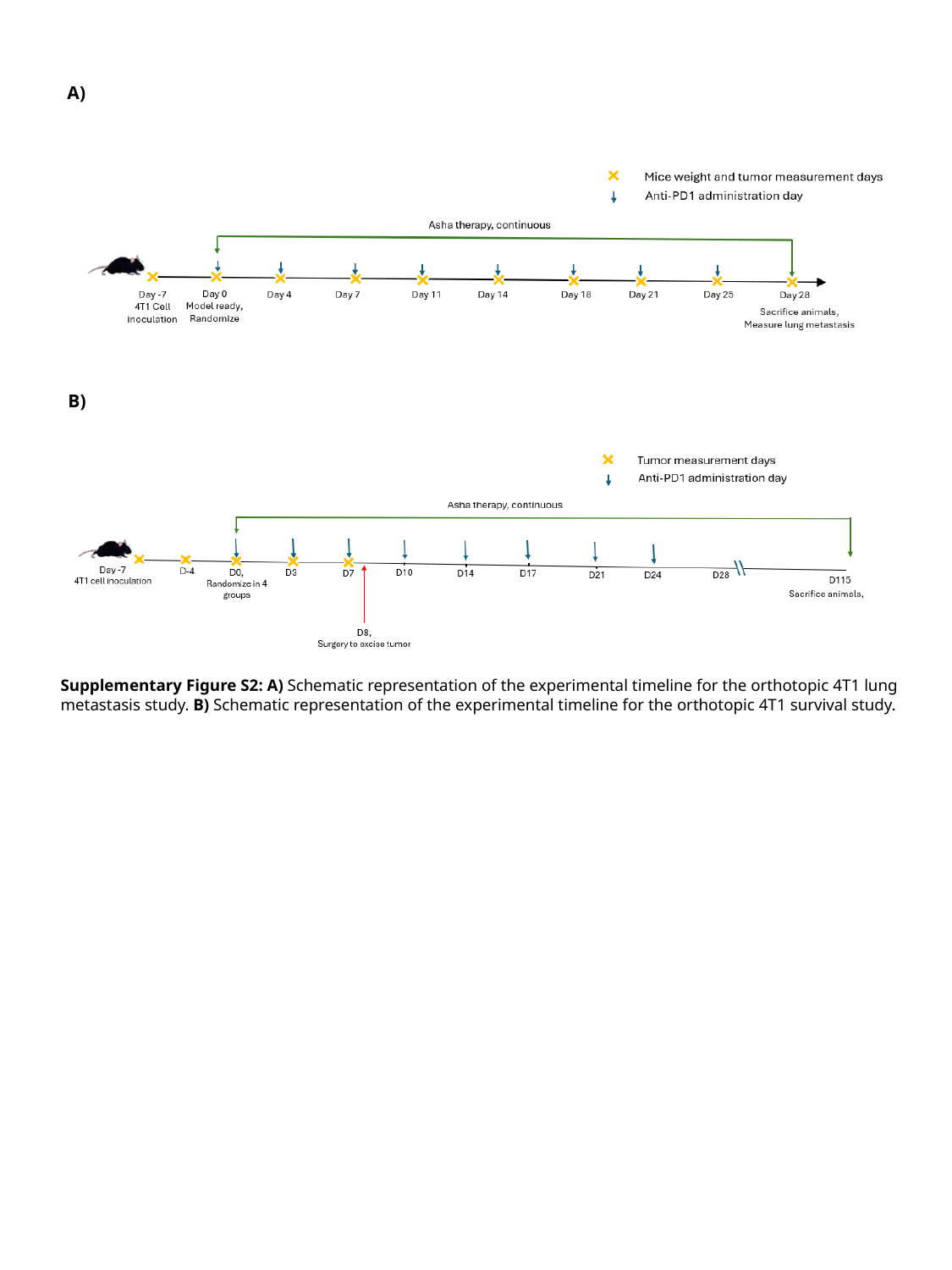

A)
B)
Supplementary Figure S2: A) Schematic representation of the experimental timeline for the orthotopic 4T1 lung metastasis study. B) Schematic representation of the experimental timeline for the orthotopic 4T1 survival study.

### Slide 3
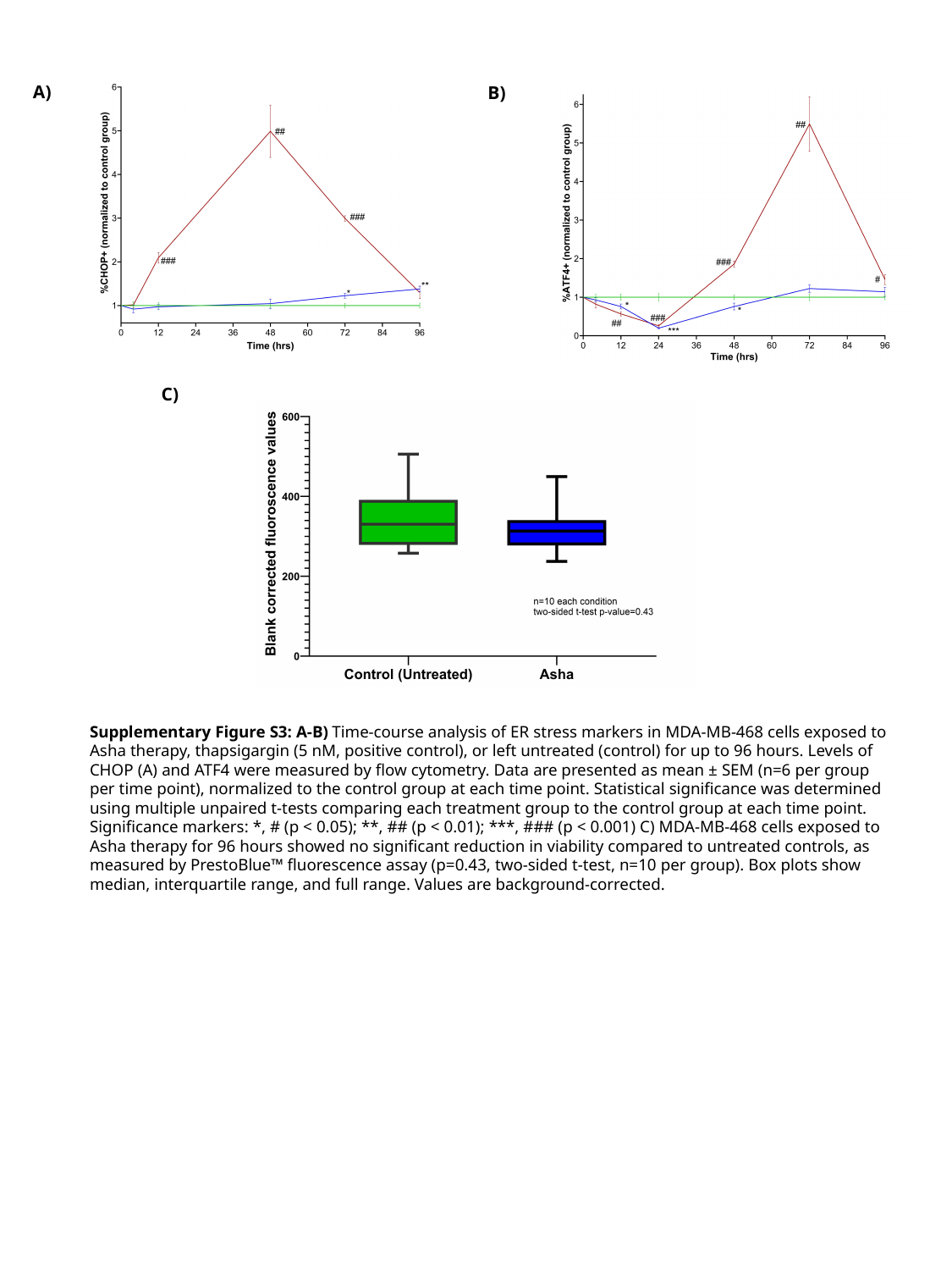

A)
B)
C)
Supplementary Figure S3: A-B) Time-course analysis of ER stress markers in MDA-MB-468 cells exposed to Asha therapy, thapsigargin (5 nM, positive control), or left untreated (control) for up to 96 hours. Levels of CHOP (A) and ATF4 were measured by flow cytometry. Data are presented as mean ± SEM (n=6 per group per time point), normalized to the control group at each time point. Statistical significance was determined using multiple unpaired t-tests comparing each treatment group to the control group at each time point. Significance markers: *, # (p < 0.05); **, ## (p < 0.01); ***, ### (p < 0.001) C) MDA-MB-468 cells exposed to Asha therapy for 96 hours showed no significant reduction in viability compared to untreated controls, as measured by PrestoBlue™ fluorescence assay (p=0.43, two-sided t-test, n=10 per group). Box plots show median, interquartile range, and full range. Values are background-corrected.

### Slide 4
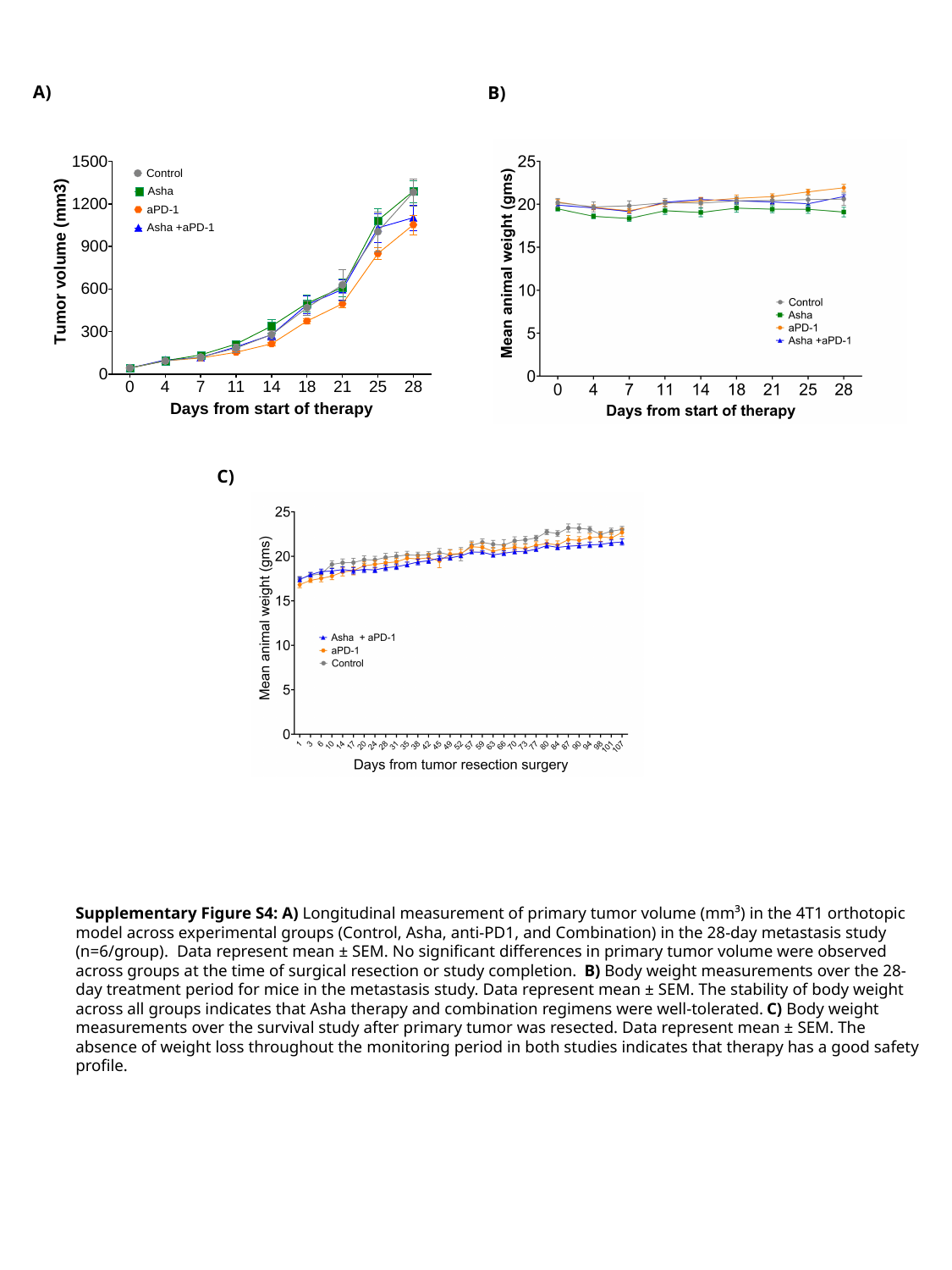

A)
B)
C)
Supplementary Figure S4: A) Longitudinal measurement of primary tumor volume (mm³) in the 4T1 orthotopic model across experimental groups (Control, Asha, anti-PD1, and Combination) in the 28-day metastasis study (n=6/group). Data represent mean ± SEM. No significant differences in primary tumor volume were observed across groups at the time of surgical resection or study completion. B) Body weight measurements over the 28-day treatment period for mice in the metastasis study. Data represent mean ± SEM. The stability of body weight across all groups indicates that Asha therapy and combination regimens were well-tolerated. C) Body weight measurements over the survival study after primary tumor was resected. Data represent mean ± SEM. The absence of weight loss throughout the monitoring period in both studies indicates that therapy has a good safety profile.

### Slide 5
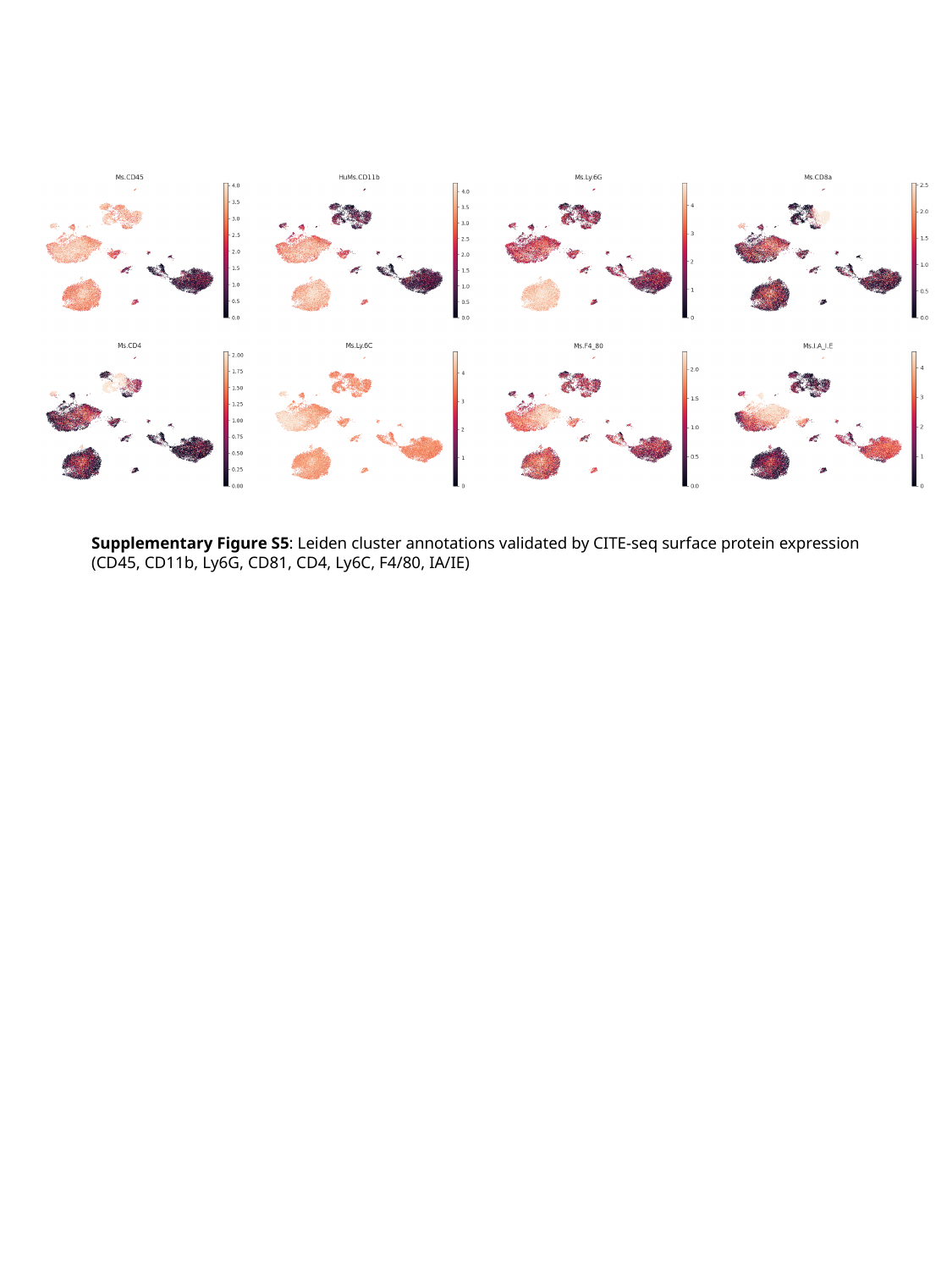

Supplementary Figure S5: Leiden cluster annotations validated by CITE-seq surface protein expression (CD45, CD11b, Ly6G, CD81, CD4, Ly6C, F4/80, IA/IE)

### Slide 6
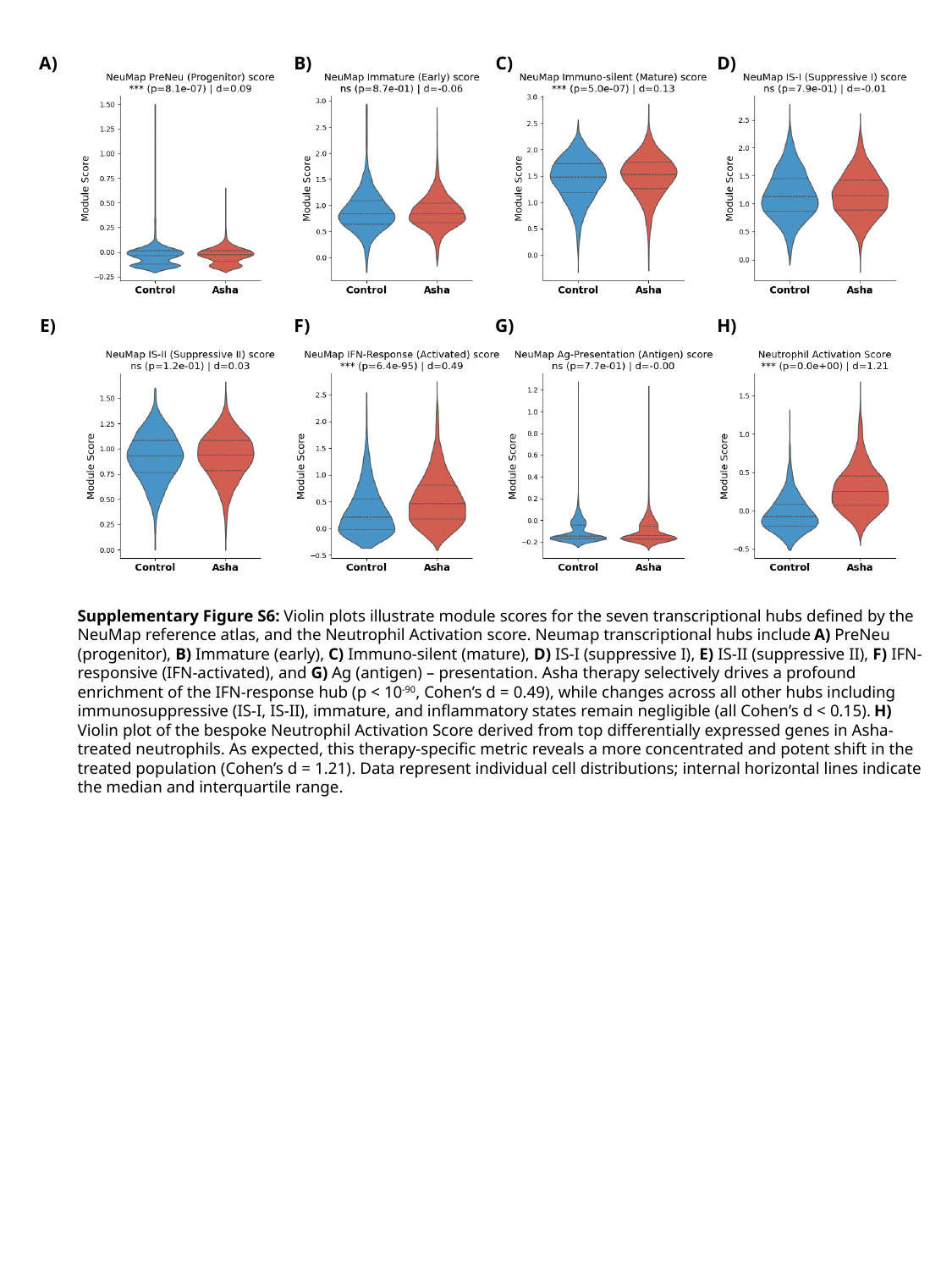

B)
A)
C)
D)
F)
E)
G)
H)
Supplementary Figure S6: Violin plots illustrate module scores for the seven transcriptional hubs defined by the NeuMap reference atlas, and the Neutrophil Activation score. Neumap transcriptional hubs include A) PreNeu (progenitor), B) Immature (early), C) Immuno-silent (mature), D) IS-I (suppressive I), E) IS-II (suppressive II), F) IFN-responsive (IFN-activated), and G) Ag (antigen) – presentation. Asha therapy selectively drives a profound enrichment of the IFN-response hub (p < 10^-90^, Cohen’s d = 0.49), while changes across all other hubs including immunosuppressive (IS-I, IS-II), immature, and inflammatory states remain negligible (all Cohen’s d < 0.15). H) Violin plot of the bespoke Neutrophil Activation Score derived from top differentially expressed genes in Asha-treated neutrophils. As expected, this therapy-specific metric reveals a more concentrated and potent shift in the treated population (Cohen’s d = 1.21). Data represent individual cell distributions; internal horizontal lines indicate the median and interquartile range.

### Slide 7
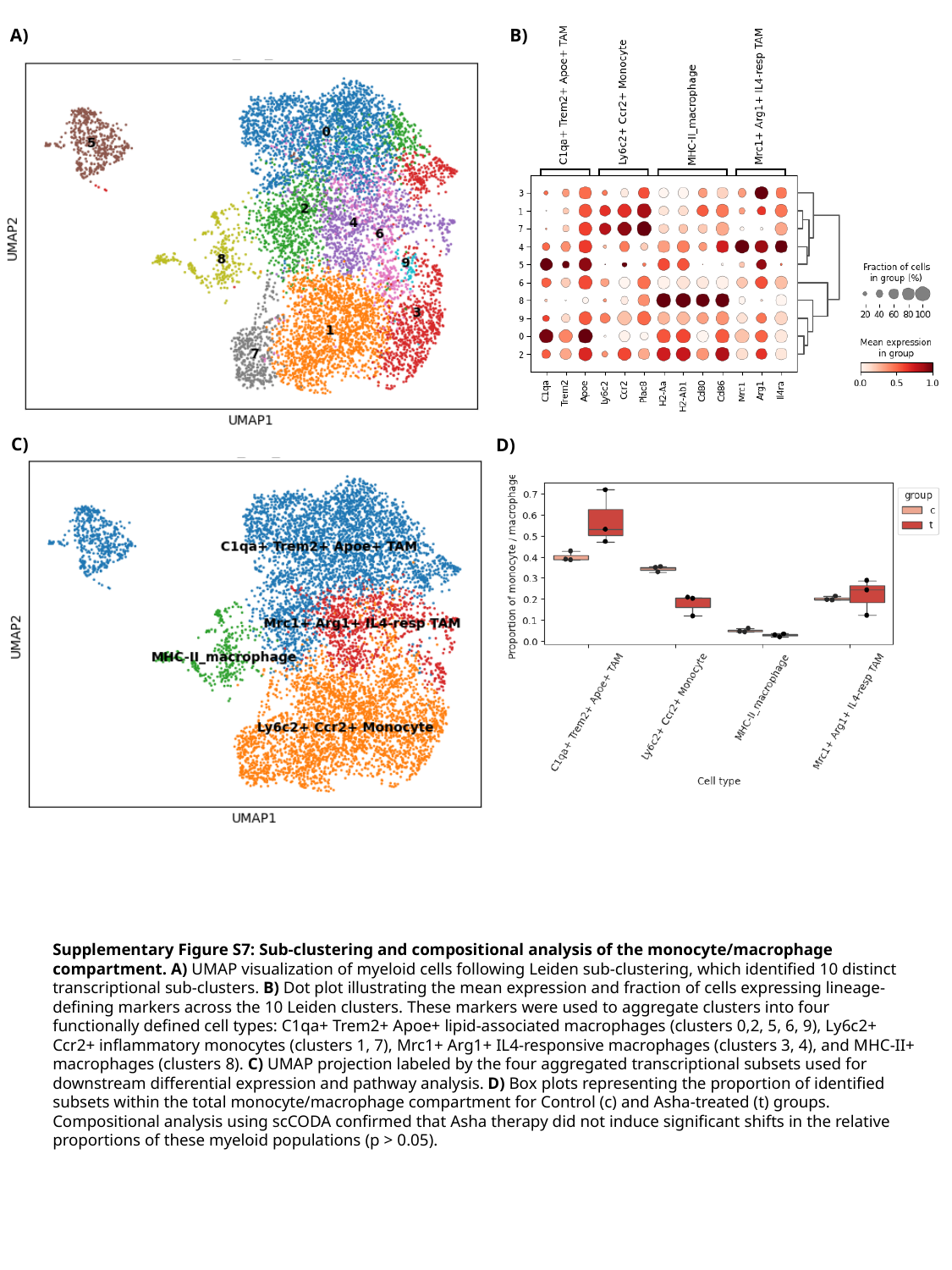

B)
A)
C)
D)
Supplementary Figure S7: Sub-clustering and compositional analysis of the monocyte/macrophage compartment. A) UMAP visualization of myeloid cells following Leiden sub-clustering, which identified 10 distinct transcriptional sub-clusters. B) Dot plot illustrating the mean expression and fraction of cells expressing lineage-defining markers across the 10 Leiden clusters. These markers were used to aggregate clusters into four functionally defined cell types: C1qa+ Trem2+ Apoe+ lipid-associated macrophages (clusters 0,2, 5, 6, 9), Ly6c2+ Ccr2+ inflammatory monocytes (clusters 1, 7), Mrc1+ Arg1+ IL4-responsive macrophages (clusters 3, 4), and MHC-II+ macrophages (clusters 8). C) UMAP projection labeled by the four aggregated transcriptional subsets used for downstream differential expression and pathway analysis. D) Box plots representing the proportion of identified subsets within the total monocyte/macrophage compartment for Control (c) and Asha-treated (t) groups. Compositional analysis using scCODA confirmed that Asha therapy did not induce significant shifts in the relative proportions of these myeloid populations (p > 0.05).

### Slide 8
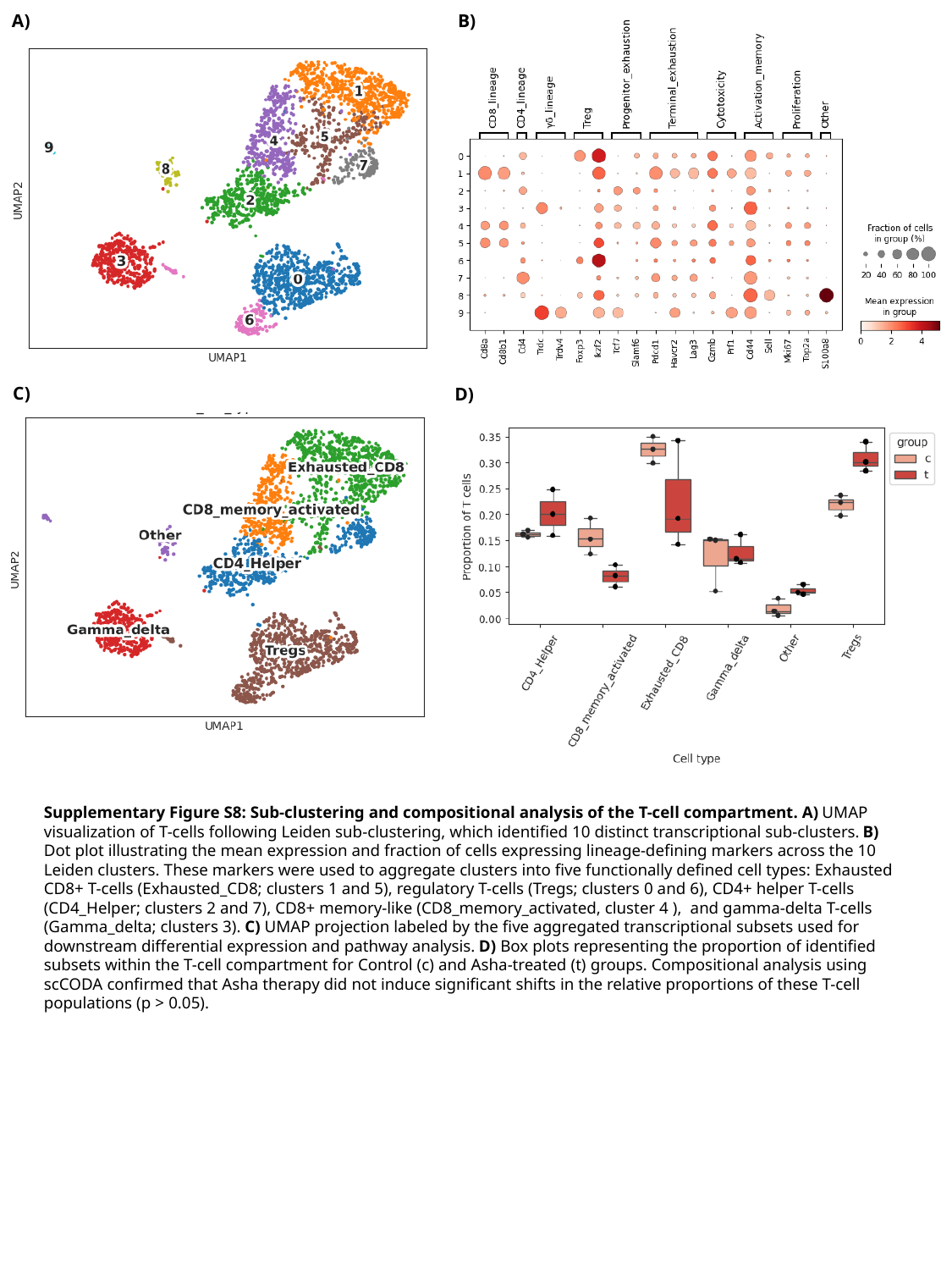

B)
A)
C)
D)
Supplementary Figure S8: Sub-clustering and compositional analysis of the T-cell compartment. A) UMAP visualization of T-cells following Leiden sub-clustering, which identified 10 distinct transcriptional sub-clusters. B) Dot plot illustrating the mean expression and fraction of cells expressing lineage-defining markers across the 10 Leiden clusters. These markers were used to aggregate clusters into five functionally defined cell types: Exhausted CD8+ T-cells (Exhausted_CD8; clusters 1 and 5), regulatory T-cells (Tregs; clusters 0 and 6), CD4+ helper T-cells (CD4_Helper; clusters 2 and 7), CD8+ memory-like (CD8_memory_activated, cluster 4 ), and gamma-delta T-cells (Gamma_delta; clusters 3). C) UMAP projection labeled by the five aggregated transcriptional subsets used for downstream differential expression and pathway analysis. D) Box plots representing the proportion of identified subsets within the T-cell compartment for Control (c) and Asha-treated (t) groups. Compositional analysis using scCODA confirmed that Asha therapy did not induce significant shifts in the relative proportions of these T-cell populations (p > 0.05).
