## Supplementary for "Magnetic field-induced ER stress reprograms the tumor microenvironment to improve triple-negative breast cancer survival"

**Supplementary 1:**

To evaluate whether the neutrophil program induced by Asha therapy is reflected in human breast cancer datasets, we derived a neutrophil interferon-stimulated gene (ISG) signature from differential expression (DESeq2) results obtained in vitro. This section details the stepwise procedure used to generate this signature, the rationale for gene inclusion and exclusion, and sensitivity analyses performed using alternative gene-set sizes (top-50 and top-200 upregulated genes).

1. **Differential expression analysis and gene ranking**

Bulk RNA-seq was performed on mouse neutrophils isolated from treated versus control tumors. Differential expression analysis using DESeq2 identified 1,114 significantly upregulated genes (padj < 0.05). To derive a robust, biologically interpretable gene set, we ranked all upregulated genes by Wald statistic, which prioritizes genes with strong effect size and low variance.

The top 100 upregulated genes by Wald statistic are shown below (mouse symbols):

*Gm42418, Cmss1, AY036118, Rps28, Ly6e, Rpl13a, Rps29, Rpl37a, Rps23, Filip1l, Tbrg1, Rpl38, Rpl36, Gm16337, Rps26, Rpl26, Gm36043, Rplp1, Rpl13, Rpl27, Rps27, Rps15, Rplp2, Lgals1, Rpl37, Rpl34, Rps21, Rpl24, Spp1, Atp5mpl, 1600014C10Rik, Psme1, Rbm8a, Rps6, Rpl23a, Rps19, Mndal, Gm10076, H2-T22, Rpl11, Sec61g, Rpl31, Manf, Rpl22, Rps14, Rpl17, Rpl39, Atp5d, Uba52, Psmb1, Gm16894, Oasl2, Xaf1, Ifit3b, Ifi27l2a, Pkig, Gm4876, Ifit3, Naca, Npc2, Rpl30, Eif2s2, Nf1, Rpl6, Gm37240, Tomm20, Gm20536, Zbp1, Rps15a, Zfas1, Rpl27a, Atp5md, Mif, Mrpl54, Map1b, Psmb8, Rps11, Ndufb9, Rpl19, Mfsd4a, Id3, Rpl15, 2410006H16Rik, Ifit2, Yrdc, 4930453N24Rik, Rpl7, Ifi211, Ndufb1-ps, Uqcrq, Arg1, Cnih4, Rpl18, Rps13, Cstdc4, Ufc1, Atp5e, B2m, Rpl12, Uqcr10*

This list includes multiple categories of genes:

- canonical interferon-stimulated genes (e.g., *Ly6e, Ifit3, Ifit2, Ifi27, Oasl2, Xaf1, Zbp1*)
- antigen-processing machinery (*Psmb1, Psmb8, Psme1, B2m*)
- ER stress–associated genes (*Sec61g, Tbrg1*)
- neutrophil-associated activation markers (*Spp1, Arg1, Mndal*)
- translational and mitochondrial housekeeping transcripts (e.g., ribosomal proteins, *Atp5* family)
- mouse-specific non-coding loci (Gm* genes)

2. **Biological and technical filtering strategy**

To build a biologically interpretable, neutrophil-specific signature appropriate for projection onto human bulk RNA-seq cohorts, we applied the following filtering steps:

1. Removed ribosomal, mitochondrial, and translation-associated genes, which represented generic metabolic responses rather than neutrophil-intrinsic immunologic programming.
2. Removed mouse-specific Gm loci lacking human orthologs (e.g., *Gm42418, Gm10076, Gm13684*), and mapped *Gm16337* to its closest human antiviral ortholog IFNL3.
3. Excluded genes not associated with neutrophil or interferon biology, including RNA-processing factors (e.g., *Rbmx, Rbm8a, U2af1, Eif2s2*) and endoplasmic reticulum housekeeping genes without known immune relevance.
4. Excluded genes whose expression in bulk tumors is dominated by non-neutrophil cell types, such as (*Arg1*, TAMs; *Lgals1*, fibroblasts; *Cxcl9*, T-cell/endothelial), to avoid confounding.
3. **Final neutrophil ISG signature gene list**

After applying these criteria, we curated a 19-gene "Asha Neutrophil ISG Signature", consisting of genes meeting both:

a) high statistical induction in Asha-treated neutrophils, and
b) clear mechanistic relevance to interferon signaling, antigen processing, and neutrophil activation.

The gene list included following genes:

LY6E, IFIT3, IFIT2, IFI27, PSMB8, SPP1, HLA-E, OASL, XAF1, ZBP1, PSMB1, PSME1, B2M, TBRG1, FILIP1L, SEC61G, CMSS1, MNDAL, IFNL3
